# Cryo-EM reveals the complex architecture of dynactin’s shoulder and pointed end

**DOI:** 10.1101/2020.07.16.206359

**Authors:** Clinton K. Lau, Francis J. O’Reilly, Balaji Santhanam, Samuel E. Lacey, Juri Rappsilber, Andrew P. Carter

**Affiliations:** Structural Studies Division, MRC Laboratory of Molecular Biology, Cambridge, United Kingdom; Bioanalytics, Institute of Biotechnology, Technische Universität Berlin, Berlin, Germany

## Abstract

Dynactin is a 1.1 MDa complex that activates the molecular motor, dynein, for ultra-processive transport along microtubules. In order to do this it forms a tripartite complex with dynein and a coiled-coil adaptor. Dynactin consists of an actin-related filament whose length is defined by its flexible shoulder domain. Despite previous cryo-EM structures, the molecular architecture of the shoulder and pointed end of the filament is still poorly understood due to the lack of high-resolution information in these regions. Here we combine multiple cryo-EM datasets and define precise masking strategies for particle signal subtraction and 3D classification. This overcomes domain flexibility and results in high resolution maps into which we can build the shoulder and pointed end. The unique architecture of the shoulder positions the four identical p50 subunits in different conformations to bind dynactin’s filament and securely houses the p150 subunit. The pointed end map allows us to build the first structure of p62, and reveals the molecular basis for cargo adaptor binding to different sites at the pointed end.

## Introduction

Dynactin is a large, multi-subunit co-activator of the molecular motor, cytoplasmic dynein 1. It is required for long-range transport along microtubules in many animals and fungi (1). Dynein-dynactin complexes transport a wide range of cargos, from large organelles such as mitochondria to small mRNA particles. Dynactin is built around an Arp1/actin filament, which is capped by pointed and barbed end complexes (2–6) (Figure 1A). On the side of the filament sits a shoulder domain from which the 75 nm-long p150Glued (DCTN1, hereafter referred to as p150) projection extends (2, 6). Dynactin binds dynein in the presence of coiled-coil cargo adaptors, such as BICD2, BICDR1 and Hook3 to form a highly processive motor complex (7, 8). Dynein contacts dynactin’s Arp1 filament via its heavy chain (6, 9), and the p150 N terminus via its intermediate chain (10, 11). Coiled-coil cargo adaptors make interactions along dynactin’s filament and pointed end and bind to dynein’s heavy chain and light intermediate chain (6, 9, 12–14).

**Figure 1.**
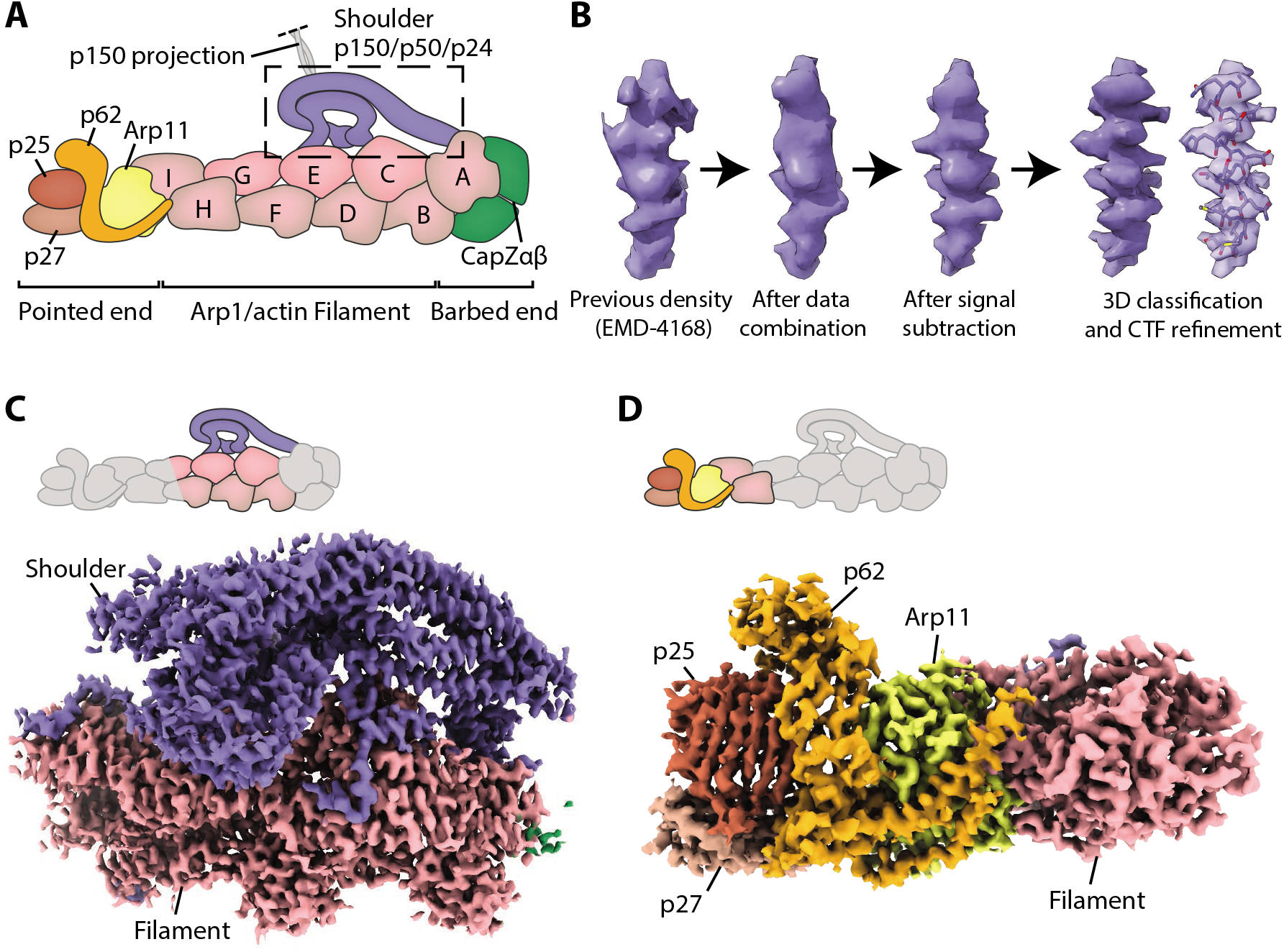
High resolution maps of dynactins shoulder and pointed end. **A.** Schematic showing the domain architecture of dynactin. **B.** Density improvements during processing. For each step, sample density is taken from the same p50 arm helix in the shoulder. **C.** Map of the shoulder region. Density of the shoulder is colored in blue. **D.** Map of the pointed end including Arp 11 (yellow); p62 (orange); p25 (brown) and p27 (light brown).

Despite multiple structures containing dynactin (6, 9), neither the shoulder nor the pointed end have been resolved at high resolution, due to flexibility between these domains and the filament. The shoulder consists of the C termini of the p150 projection, four copies of p50 (DCTN2) and two p24 (DCTN3) subunits (3). The N termini of the p50 subunits bind the Arp1 filament and act as molecular rulers to determine its length (6, 15, 16). Previous studies showed that the shoulder contains long three-helical bundles with a two-fold pseudo-symmetry (6). However due to the limited resolution in the shoulder, it was not possible to assign individual subunits. Key outstanding questions include how the C termini of p150 are embedded into the shoulder, how the four p50 subunits organize into a structure with two-fold symmetry, and how their N termini project to correctly bind the filament.

The pointed end is important for binding dynein-dynactin cargo adaptors (6, 9, 13, 17–19). It consists of four subunits: actin-related protein 11 (Arp11, ACTR10); p62 (DCTN4); and the closely related p25 and p27 (DCTN5 and DCTN6). Previous maps were sufficient to build Arp11 and place, but not assign, models of p25 and p27 (6, 9, 20). It was not possible to model the structure of p62 due to poor density and lack of structural homologs. Using the previous dynein tail-dynactin-BICD2 structure (6), structural modelling with molecular dynamics predicted residues in p25 to bind to all coiled-coil cargo adaptors (21). However, subsequent structures revealed that Hook3 and BICDR1 in fact contact different regions of the pointed end (9). The lack of high resolution in these regions means that it is currently unclear which pointed end residues interact with the different cargo adaptors.

To overcome the flexibility within dynactin, we combined multiple cryo-EM datasets of different dynactin-containing complexes and developed a precise masking strategy for signal subtraction. In combination with other recent advances in cryo-EM data processing, this allowed us to produce a 3.8 Å map of dynactin’s shoulder and a 4.1 Å map of the pointed end. We find that the p150 C termini are securely anchored into the shoulder by making extensive interactions with other subunits. The p50 subunits are asymmetrically arranged in four unique conformations to position their N termini correctly to bind to dynactin’s filament. At the pointed end of dynactin, we build an atomic model of p62 and identify the residues involved with cargo adaptor binding. We also resolve the pointed end residues that interact with the p150 projection when it folds back to contact dynactin. We find that in this conformation p150 overlaps with all adaptor binding sites, suggesting that it acts to inhibit dynactin’s interactions with cargo adaptors.

## Results

### Determination of high resolution structures of dynactin’s shoulder and pointed end

One cause of limited resolution in previous dynactin structures was flexibility that smeared the density in peripheral regions. Particle signal subtraction can overcome this by computationally subtracting density around regions of a protein complex that move as a rigid body, allowing further refinement of the rigid body to higher resolution (22). In our previous structure of dynein tail-dynactin-BICDR1 (TDR), this allowed us to build an atomic model of the dynein tails (9). Using that dataset, we first attempted to implement the same strategy for dynactin’s shoulder and pointed end. However signal subtraction on these regions using the TDR dataset alone did not produce maps of sufficient quality to build an atomic model. To overcome this, we decided to increase our particle number, then use signal subtraction with mask optimization, 3D classification, CTF refinement and particle re-centering to increase the resolution.

We increased our dataset size by combining data from our previous dynactin (6), TDR and dynein tail-dynactin-Hook3 (TDH) structures (9), and by incorporating new TDH data (Supplementary Figure 1, Table 1). For the TDR and TDH datasets we used signal subtraction to remove density for the dyneins and cargo adaptors. Using the resulting dynactin par-ticles we focused on either the shoulder or pointed end, and performed signal subtraction and local refinement for each region (Figure 1B, Supplementary Figure 1). To determine the best possible mask for this process, we first tested a broad range of masks to identify regions that could be refined to higher resolution. We proceeded with masks that gave the maps containing the best density.

**Table 1:** Electron microscopy data collection and refinement statistics

|  | Dynein tail-Dynactin-Hook3 | Dynein tail-Dynactin-Hook3<sub>Leeds</sub> | Dynactin<sub>p150docked</sub> |
| --- | --- | --- | --- |
| **Data collection and processing** |  |  |  |
| Voltage (kV) | 300 | 300 | 300 |
| Electron exposure (e<sup>-</sup>/Å<sup>2</sup>) | 40 | 40 | 40 |
| Pixel size (Å) | 0.58 | 1.07 | 1.16 |
| Micrographs | 2136 | 1263 | 26,036 |
| Symmetry imposed | C1 | C1 | C1 |
| Initial particle images (no.) | 77,941 | 34,276 | 1,361,241 |
| Final particle images (no.) | 13,799 | 4,567 | 98,511/6,342<sup>1</sup> |

|  | Overall | Shoulder | Pointed End | Pointed<br>end:BICDR1 | Pointed<br>end:Hook3 | Pointed<br>end:p150 | Pointed<br>end:BICD2 |
| --- | --- | --- | --- | --- | --- | --- | --- |
| **Reconstruction** |  |  |  |  |  |  |  |
| Map | EMD-11313 | EMD-11314 | EMD-11315 | EMD-11317 | EMD-11318 | EMD-11319 | EMD-2860<sup>2</sup> |
| Particle number | 336,972 | 103,532 | 132,845 | 205,611 | 57,348 | 20,037 |  |
| Map resolution (Å) | 3.8 | 3.8 | 4.1 | 4.1 | 4.5 | 6.8 |  |
| FSC threshold | 0.143 | 0.143 | 0.143 | 0.143 | 0.143 | 0.143 |  |
| Accuracy |  |  |  |  |  |  |  |
| Rotation (°) | 1.10 | 1.11 | 2.64 | 2.64 | 1.87 | 3.74 |  |
| Translation (Å) | 1.14 | 0.91 | 1.18 | 1.39 | 1.22 | 2.18 |  |
| Map sharpening |  |  |  |  |  |  |  |
| B factor (Å<sup>2</sup>) | -90 | -118 | -167 | -70 | -100 | -100 |  |
| Refinement |  |  |  |  |  |  |  |
| Map CC (around atoms) | 0.70 | 0.74 | 0.75 | 0.75 | 0.72 | 0.68 | 0.80 |
| Model composition |  |  |  |  |  |  |  |
| PDB | 6ZNL | 6ZNL | 6ZNL | 6ZNM | 6ZNN | 6ZNO | 6ZO4 |
| Non-hydrogen atoms | 53,001 | 24,799 | 14,481 | 19,256 | 19,460 | 19,740 | 19,305 |
| Protein residues | 7,016 | 3,409 | 1,899 | 2,470 | 2,320 | 2,320 | 2,320 |
| Ligands (ADP/ATP/Zn) | 9/1/3 | 4/0/0 | 2/1/3 | 2/1/3 | 2/1/3 | 2/1/3 | 2/1/3 |
| R.m.s. deviations |  |  |  |  |  |  |  |
| Bond lengths (Å) | 0.01 | 0.01 | 0.01 | 0.01 | 0.01 | 0.01 | 0.014 |
| Bond angles (°) | 1.79 | 1.82 | 1.76 | 1.72 | 1.73 | 1.70 | 1.72 |
| Validation |  |  |  |  |  |  |  |
| MolProbity score | 1.45 | 1.37 | 1.17 | 1.39 | 1.40 | 1.39 | 1.41 |
| Clashscore | 2.73 | 2.36 | 0.55 | 1.79 | 1.73 | 1.66 | 1.82 |
| Poor rotamers (%) | 0.02 | 0.04 | 0.00 | 0.00 | 0.00 | 0.00 | 0.00 |
| Ramachandran plot |  |  |  |  |  |  |  |
| Favored (%) | 94.25 | 94.89 | 92.03 | 92.92 | 92.47 | 92.47 | 92.47 |
| Disallowed (%) | 0.04 | 0.08 | 0.00 | 0.00 | 0.00 | 0.00 | 0.00 |
| Cβ deviations (%) | 0.05 | 0.06 | 0.00 | 0.00 | 0.00 | 0.00 | 0.00 |

<sup>1</sup> Number of particles containing p150 docked onto filament
<sup>2</sup> (Urnavicius et al. 2015)

For the shoulder, we next further optimized the mask. This was accomplished by testing the mask using focused refinement without signal subtraction. We examined the boundaries of the output map to identify further density to include or exclude (Supplementary Figure 2). Specifically, we adjusted the shoulder mask to exclude Arp1 subunit A, which was slightly flexible relative to the rest of the map, and include parts of Arp1 subunits F and G, which were well-resolved, but outside our original mask. Then we used the optimized mask for signal subtraction. This resulted in better density throughout the map compared to non-optimized versions, allowing us to build a more complete structure. CTF refinement followed by 3D classification for the shoulder region further improved the density (Figure 1C, Supplementary Figure 1, Supplementary Figure 3).

For the pointed end, we used a relatively large mask for initial signal subtraction. We simultaneously re-centered our particles, using the new feature in RELION 3.1 (23), as this region is located far from the center of the particle. Recentering permits more meaningful priors for refinement and allowed the visualization of β-strands in p62 for the first time. After the first round of subtraction, the mask was further optimized before a second round of signal subtraction using the same strategy as described above. This was followed by 3D classification and further refinement (Figure 1D, Supplementary Figure 1, Supplementary Figure 4).

Using the new maps we could build models for the shoulder and pointed end (Figure 2, 3). To validate our structures, we crosslinked dynactin with bis(sulfosuccinmidyl)suberate (BS3) and identified the crosslinked residues using crosslinking mass spectrometry. 527 crosslinks were identified with a false discovery rate of 2% and were compared against our structure (Supplementary Figure 5A). 246 of the crosslinked residues pairs are in structurally ordered regions in dynactin. A further 59 pairs have one or both residues contained in short disordered loops that can be modelled. These 305 crosslinks satisfy the maximum theoretical length of BS3, 30 Å (C_*α*_-C_*α*_) (Supplementary Figure 5B). Of the 18 overlength crosslinks, 7 can be explained by minor structural flexibility. The other 11 are incompatible with our structure, consistent with our false discovery rate. The remaining crosslinks use at least one residue in long disordered loops (35 crosslinks) or within the p150 projection (169 crosslinks).

**Figure 2.**
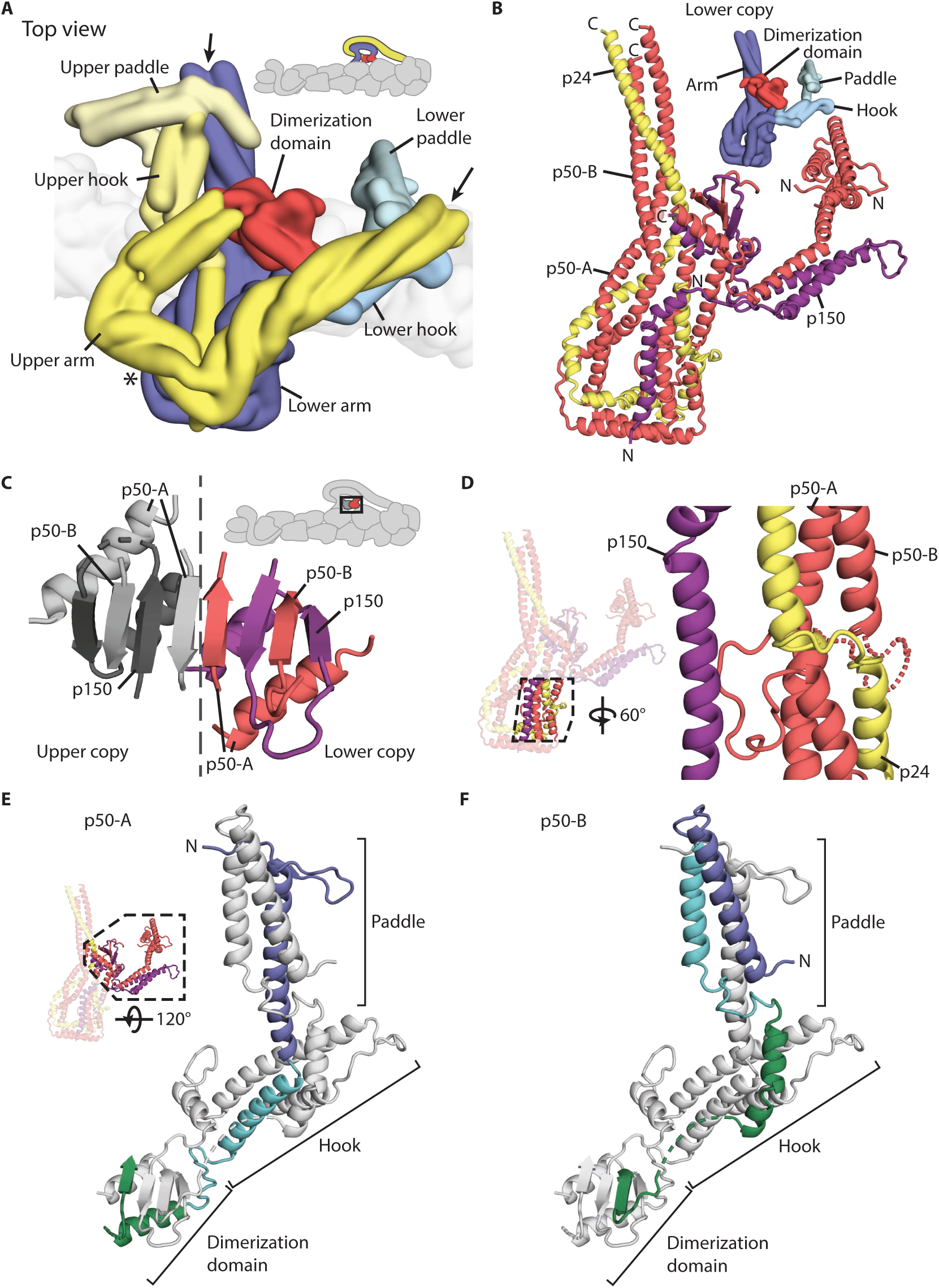
Structure of dynactins shoulder. **A.** Surface rendering showing the arrangement of the upper copy (yellows) and the lower copy (blues) in dynactins shoulder. The subdomains are shown in different color shades, and the dimerization domain is colored red. Density for the arrowed region was worse in the lower copy than the upper copy. Side chain density was absent in both copies in the middle of the arms (asterisk). **B.** The structure of the lower copy from the shoulder showing the organization of p150 (purple), p24 (yellow) and the two copies of p50 (both red). The N and C termini of each chain are shown. **C.** The dimerization domain of the shoulder. Subunits from the lower copy are colored, with the p50 subunits in red and p150 subunits in purple. **D.** Helical bundle break in the lower copy arm, showing how the three helices in the arm break and reform after a twist. p24 (yellow) forms a short loop at the break, whereas p50 (red) forms a longer loop, which is ordered in p50-A, and disordered in p50-B, modelled by a dotted line. **E,F.** The N-terminal halves of p50-A (E) and p50-B (F) adopt different conformations in each asymmetric copy (lower copy shown). In both diagrams the polypeptide is colored blue, cyan, then green to show equivalent structural features.

**Figure 3.**
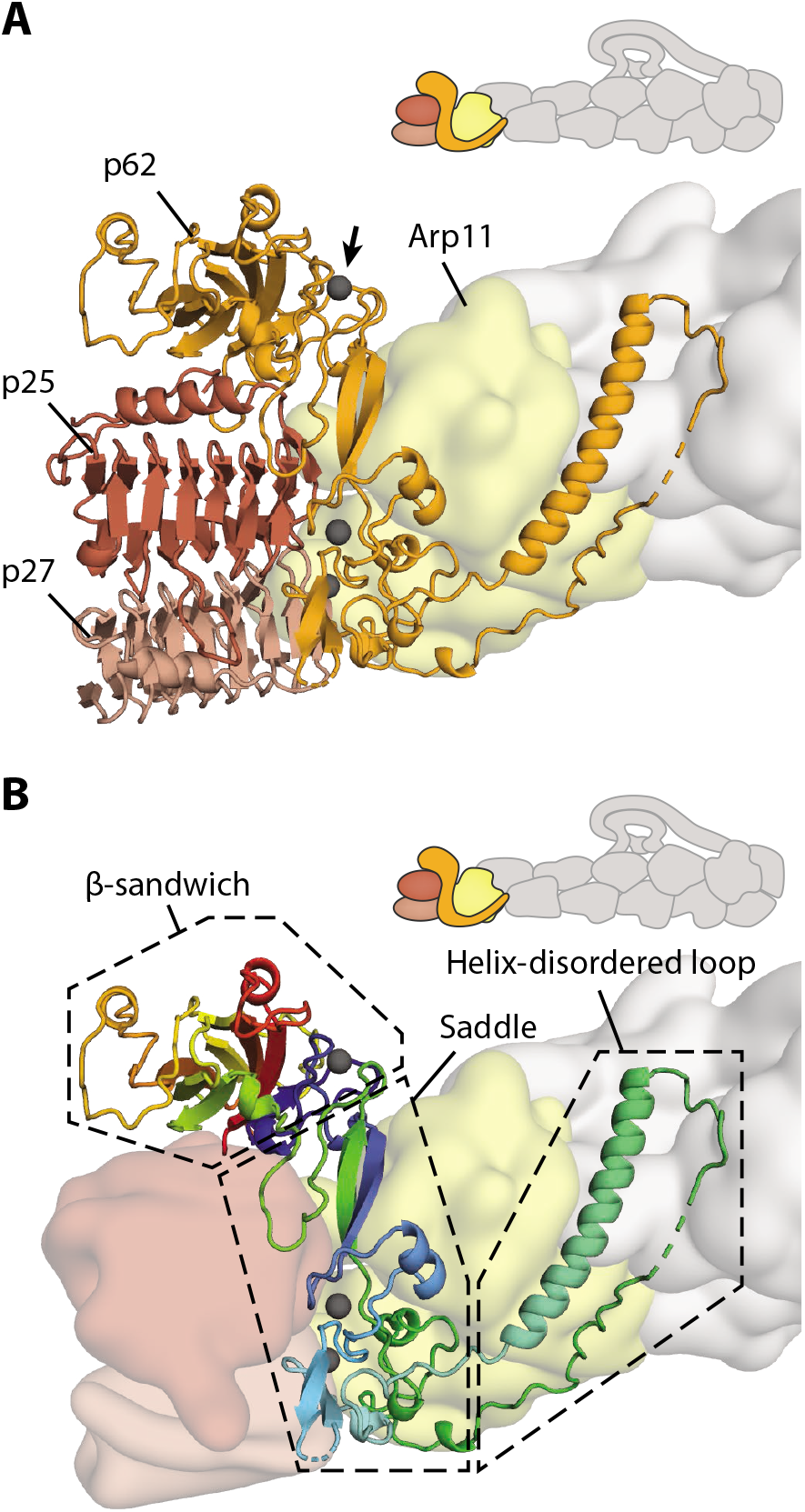
The organization of the pointed end. **A.** The structure of the pointed end showing p62 (orange), p25 (brown) and p27 (light brown) in cartoon representation. Arp11 (yellow) and the filament are shown in surface representation. Metal ions in p62 are colored gray, with the metal ion coordinating the N and C termini of p62 highlighted (arrow). **B.** Detailed structure of p62 (cartoon), colored from N to C termini in blue, cyan, green, yellow, orange then red. Dotted boxes show the different structural features of p62.

### The complex architecture of dynactin’s shoulder

Dynactin’s shoulder can be split into two halves that contain the same structural elements but are in different conformations (Figure 2A). The two halves stack on top of each other, with the lower copy making more interactions with dynactin’s filament than the upper copy. We previously observed that each copy consists of a long three-helical arm, two shorter helical bundles named the hook and the paddle, and half of a central dimerization domain (6). However, without higher resolution data, it was impossible to assign which density belonged to each subunit.

Our new map reveals density for the dimerization domain and the majority of the lower copy with sufficient resolution to build side chains. Density for the distal region of the lower arm is slightly smeared, but this region is well resolved in the upper copy (Figure 2A, arrows). The only section which has weaker side chain density in both copies is in the middle of the arms (Figure 2A, asterisk). By combining information from both copies, we can build a model for the whole shoulder. Figure 2B shows the complete structure of the lower copy containing one p150 C terminus (purple), two p50 subunits (p50A and p50B – red) and one p24 (yellow).

The p150 C terminus (residues 1090-1140) enters the shoulder and runs along the arm, as previously proposed (6). Our structure shows it forms a helical hairpin that makes up two-thirds of the hook domain (Figure 2B). The polypeptide (residues 1253-1286) then unexpectedly folds into the dimerization domain, contributing two β-strands and one α-helix (Figure 2C). The way in which p150 is intricately interwoven with other shoulder subunits strongly suggests that it is an obligate part of dynactin’s shoulder (Figure 2B). The presence of p150 in the dimerization domain poses an architectural puzzle for the identification of the surrounding secondary structural elements. Our previous structure showed this domain consists of an eight-stranded β-sheet and four α-helices, with a two-fold rotational symmetry (6). With each p150 contributing two strands to the sheet and one helix, the remaining four strands and two helices must be split between the four identical p50 subunits (Figure 2C). Our structure shows that all four p50 subunits contribute a β-strand to the dimerization domain, but only two of the four subunits add a helix.

In both halves of the shoulder, a long three-helical arm extends from the dimerization domain. Each arm consists of the C-terminal portions of both p50 subunits (residues 217-403), and the entirety of the shorter p24 (Figure 2B). Along the length of the arm, equivalent residues in the two p50 subunits run approximately alongside each other. Surprisingly as the arm extends away from the dimerization domain, the helical bundle breaks, twists by 120°, then reforms (Figure 2D). In p24 this break is spanned by a short loop, visible in our structure. In p50 there are much longer loops between the pre-break and post-break helices (Figure 2D, red lines). In one p50 copy, p50-A, this long loop is ordered (Figure 2D, solid red line), packing against the p150 subunit and hence contributing to the stability of the structure.

In contrast to the similar C-terminal portions of the p50s, the N-terminal sections adopt two remarkably different conformations in the dimerization domain, hook and paddle (Figure 2E,F). Whereas residues 100-159 in p50-A contribute one helix to each of the paddle and hook domains (Figure 2E, blue and cyan), the same sequence in p50-B forms a helical hairpin in the paddle (Figure 2F). In the dimerization domain the two p50s contribute different strands to theβ-sheet, as described above. Finally, a short helix (residues 174-182) lies behind the dimerization in p50-A (Figure 2E, green helix). The equivalent helix in p50-B is 60 Å away from the β-sheet and close to the hook domain (Figure 2F), contacting the other arm (Supplementary figure 6A). This radically different arrangement is facilitated by a long loop between the short helix and β-strand, which is completely flexible p50-A, but pulled almost taut in p50-B (Figure 2F, dotted line, Supplementary Figure 6A). The different arrangement of the two p50 subunits in our model is validated by our mass spectrometry data. Crosslinks with other dynactin proteins are satisfied by these two conformations of p50, whilst not satisfied by one conformation of p50 alone (Supplementary Figure 6B).

As a consequence of the different p50 conformations, the N-terminal 100 residues of the two p50s in each half of the shoulder project from opposite sides of the paddle domains (Figure 2E, F). Combined with the asymmetry between copies in the shoulder, this orients the extended N termini to bind dynactin’s filament at four distinct sites, in order to interact with all eight Arp1 subunits (6).

### The assembly of dynactin’s pointed end complex

In previous structures of dynactin’s pointed end, only Arp11 showed density for side chains (6). In our new map we can now build structures of p62, p25 and p27 and determine how they interact.

The related p25 and p27 both adopt similar left-handed β-helical folds (20). They have slight differences in their C-terminal helices, with p27 containing a shorter helix than p25. This, combined with side chain differences, allowed us to unambiguously assign the two proteins (Figure 3A, Supplementary Figure 7A). p62 adopts an unusual fold (Figure 3B, Supplementary Figure 8). The N-terminal and C terminal β-sheets come together to form a β-sandwich domain. The central portion of p62, which we call the saddle, contains multiple cysteines that fold into three zinc-binding motifs, with density between the cysteines for metal ions. A long helix extends from the middle of the saddle domain and is followed by a partially disordered loop.

The p62 saddle wraps around the Arp11 subunit at the end of dynactin’s filament. The long helix-loop structure folds back across the surface of Arp11 and contacts the neighboring β-actin subunit in the filament. p25 is located between the p62 β-sandwich and p27. It makes a small contact (134 Å^2^ surface area) with Arp11 (subdomain 2, Supplementary Figure 7B) but is predominantly held in place by its interactions with p62 (2746 Å^2^ SA). Its β-helical fold contacts the p62 saddle and its C-terminal helix makes interactions with p62’s β-sandwich domain. p27 binds to p25 via an extensive interface along their β-helical folds (1667 Å^2^ SA) (6). It also makes a small contact with Arp11 (subdomain 4, 263 Å^2^ SA, Supplementary Figure 7B) and a number of interactions with p62’s saddle (635 Å^2^ SA), albeit fewer than p25.

### Interaction sites for cargo adaptors on the pointed end complex

Our previous structures showed that dynein’s cargo adaptors BICDR1, BICD2 and Hook3 use overlapping, but different sites along the dynactin filament (6, 9). Here we asked which residues on dynactin’s pointed end interact with the different cargo adaptors. We performed signal subtraction on the TDR and TDH datasets individually to focus on the pointed end, which slightly improved the density around the cargo adaptor interaction sites compared with previous maps (9), particularly at the distal end. We docked our new structure into these maps, and also into our previous dynein tail-dynactin-BICD2 structure (6).

The majority of pointed end interactions cluster around four sites (Figure 4A, 4B). Site 1 involves the disordered loop following the long helix in p62 (Figure 4A). It is shared by all three adaptors, with the loop appearing to adopt different conformations to bind each adaptor. The other sites are contacted by different subsets of adaptors (Figure 4B). Site 2 is in the p62 saddle region near to p25. Site 3 is in a loop that extends out from p25, whereas site 4 is on the end face of the p25 β-helical fold. BICDR1 contacts sites 2 and 4. The main coiled coil of Hook3 contacts site 3 at the pointed end. The TDH structure revealed a second coiled coil in addition to the main one from Hook3 (9). Our structure shows that this second coiled coil binds to site 2, using a different subset of residues compared to BICDR1, and a loop in the p62 β-sandwich. BICD2 uses sites 2, 3 and 4. It interacts with another subset of residues at site 2, but the same residues at site 3 and site 4 as used by Hook3 and BICDR1 respectively.

**Figure 4.**
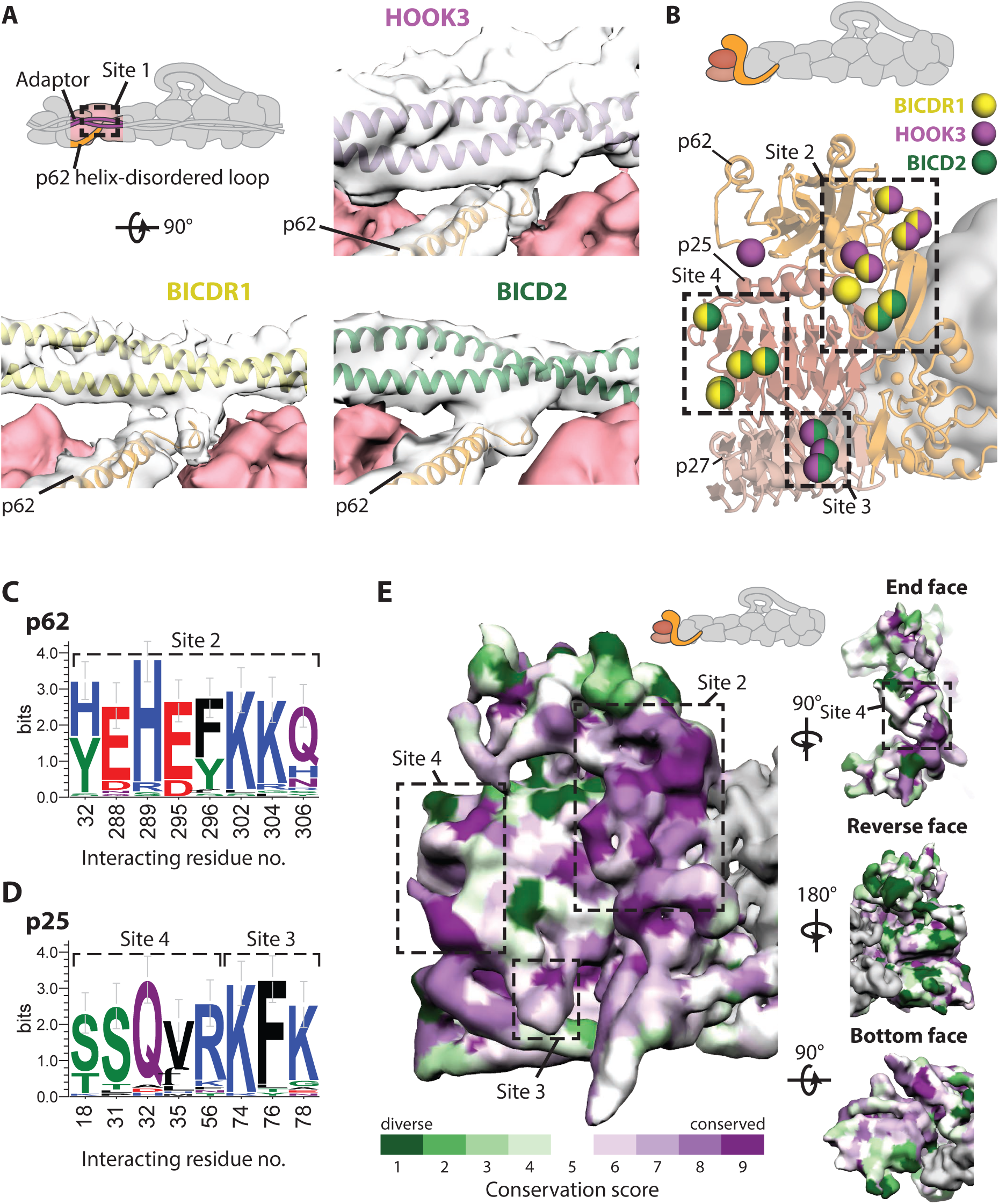
Pointed end cargo adaptor interaction sites. **A.** Site 1 in the disordered loop of p62 (orange, in clear density), binds to each cargo adaptor (BICDR1 in yellow, Hook3 in purple, BICD2 in green). Density can be seen between p62 and the adaptor at low threshold. BICDR1- and Hook3-containing maps are filtered to 6 Å. **B.** Residues on dynactins pointed end (cartoon) that interact with cargo adaptors are shown as spheres, colored according to which adaptor they bind (yellow denotes binding to BICDR1, purple to Hook3 and green to BICD2). Binding sites 2-4 are shown in boxes. **C,D.** WebLogos of the conservation of residues that interact with cargo adaptors from p62 (C)) and p25 (D) amongst diverse eukaryotes. Residues are colored by chemical property. **E.** Conservation of the surface residues plotted onto the density from the pointed end map, filtered to 6 Å. Sites 2-4 for cargo adaptor interaction are marked on the front face and end face. Conservation scores are from ConSurf, with lower conservation shown in green, and higher conservation in purple.

Different coiled-coil cargo adaptors show very limited sequence conservation (1) and, as described above, bind to different sites on the pointed end. We therefore wanted to ask if these sites are themselves conserved. We aligned sequences from a diverse set of eukaryotes that contained the pointed end proteins p62, p25 and p27 and analyzed the conservation of the residues that contact adaptors. In site 2, five of the eight residues strongly conserve their charge (E288, H289, E295, K302 and K304), and two residues (Y32 and F296) are largely aromatic (Figure 4C, upper panel). In site 3, p25 residue 74 is always positively charged and residue 76 is often aromatic (Figure 4D). In site 4 two positions (p25 residues 18 & 31) are strongly conserved as serines or threonines, and residue 32 is conserved as a glutamine (Figure 4D).

In the case of site 1, residue positions are not well conserved as the loop that contacts the cargo adaptors varies in length. Sequence analysis shows, however, that the first half of the loop maintains a net positive charge (Supplementary Figure 9). In our TDR structure we previously estimated the registry of the BICDR1 coiled coil using density for the sole tryptophan (W166) (9). This positions a series of negatively-charged glutamates near the first half of the loop in p62, suggesting an interaction between the cargo adaptor and site 1 at this point.

A plot of the surface conservation of the whole pointed end complex shows that the front side, where the cargo adaptors bind, contains several patches of strong conservation, whereas the reverse face exhibits almost none (Figure 4E). The patches of conservation overlap with sites of adaptor binding. This suggests most adaptors that bind dynactin’s pointed end interact with its front face using the sites described here.

### Dynactin p150 fold back sterically blocks all adaptors from binding

In our previous structure of dynactin alone (6), 10% of the particles showed the p150 arm folded back and docked onto the filament (Figure 5A, cartoon). The region that contacted the pointed end was assigned as two coiled coils from the N terminus of p150 called CC1A and CC1B (6, 24).

**Figure 5.**
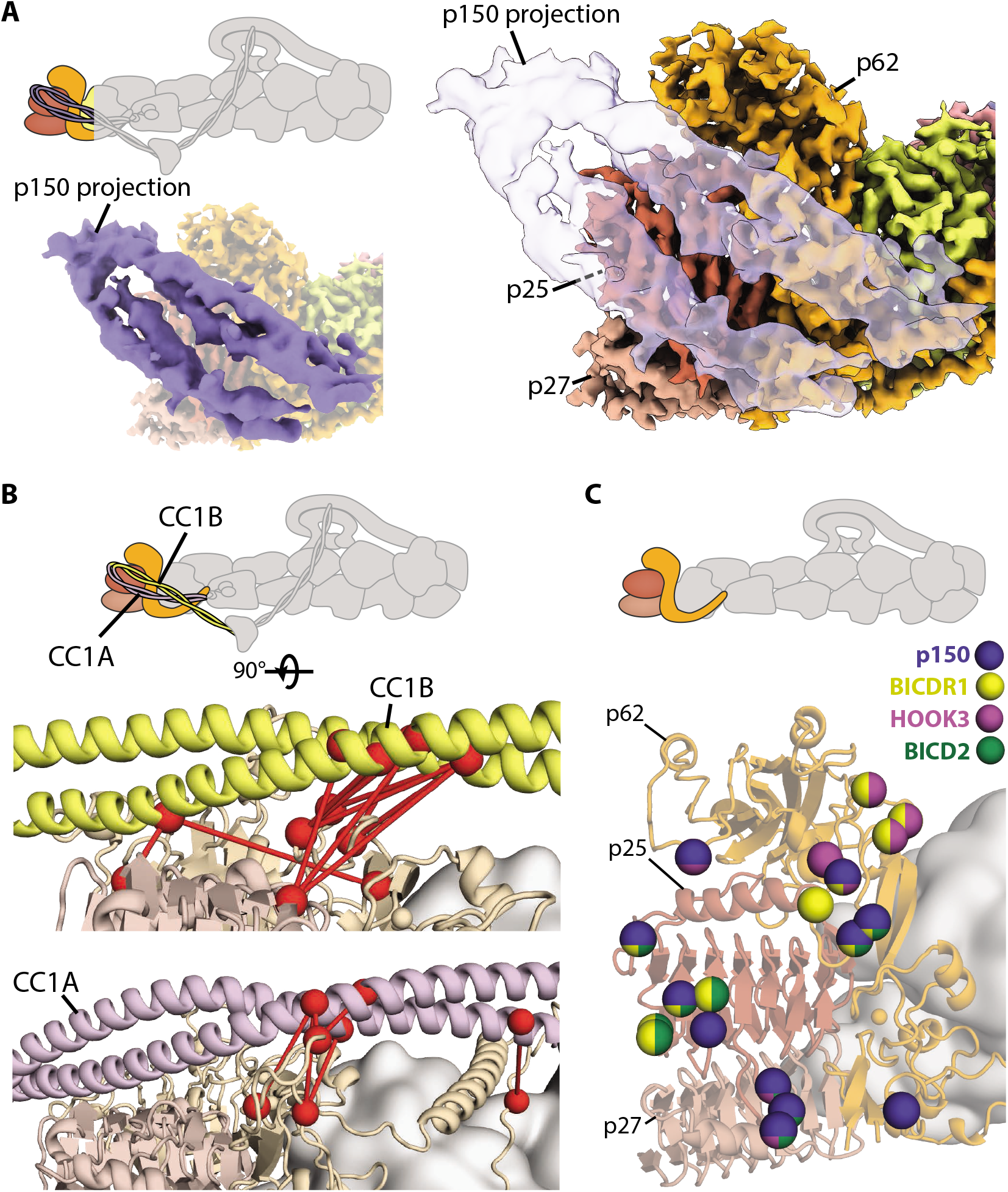
Docked p150 interaction sites overlap with adaptor binding sites. **A.** Density representation of the p150-docked structure (solid blue in left inset, transparent in right) superposed over the high resolution pointed end map shows where the coiled coil sits. **B.** Crosslinks (red dotted lines) connecting residues (shown in red spheres) between either CC1B (upper panel, in yellow) or CC1A (lower panel, in purple) and the pointed end (light orange and light brown). **C.** Residues on the pointed end that interact with the p150 or cargo adaptors are shown as spheres. Residues are colored based on their interaction partners. Those that interact with the p150 projection are shown in blue, with interactions with cargo adaptors shown in yellow (BICDR1), purple (Hook3) and green (BICD2).

Here, our crosslinking mass spectrometry data show crosslinks between CC1A, CC1B and the pointed end of dynactin (Figure 5B). This shows that the p150 docked conformation exists in solution and confirms the identity of the two coiled coils. We also find direct crosslinks from CC1A to CC1B that support their suggested anti-parallel arrangement (Supplementary Figure 10) (6, 24–26).

In p150 there is a basic domain and a small globular Cap-Gly domain, N-terminal to the CC1A/B hairpin (25). Our crosslinking shows that these regions can contact all parts of the p150 apart from the C-terminal domain, which is buried in the shoulder (Supplementary Figure 10). They are also able to interact with Arp11, actin and the Arp1 filament, consistent with them being highly mobile. In contrast, the tip of the CC1/B hairpin (residues 245-265), which contains lysine residues, makes no crosslinks to other regions. This is consistent with the suggestion that the CC1A/B hairpin, ICD and CC2 are somewhat rigid and predominantly in an extended conformation (27).

We attempted to collect more dynactin data to improve the resolution of the docked conformation. Although this only resulted in a modest improvement in resolution, our new map enabled us to better distinguish the coiled coils and interaction interface. We fit our dynactin structure into the map to examine the residues on the pointed end contacted by the p150 (Figure 5A). This shows that p150 interacts with distinct set of residues, which overlap with the sites used by cargo adaptors (Figure 5C). CC1A interacts with site 3 and site 4, whereas CC1B covers site 2. This overlap suggests when the p150 arm is bound to the pointed end, all three adaptors (BICD2, BICDR1 and Hook3) will be sterically prevented from binding to dynactin.

## Discussion

### The unique architecture of dynactin’s shoulder

Our previous structures showed that the four p50 N termini emerge from the shoulder and bind to dynactin’s filament at four distinct positions (6, 9). However, it was unclear how the p50 subunits were arranged within the shoulder to facilitate this. Our new structure reveals that each asymmetric half of the shoulder contains two p50 subunits, folded into remarkably different conformations. This arrangement correctly positions each of the four p50 N termini to bind to its cognate site on the Arp1 filament.

Despite the difference in p50 conformations, the four projecting p50 N termini are all the same length. Though it had previously been shown that residues 1-87 from p50 could bind the filament (16), it became evident from the previous dynactin structures that the four p50 binding sites on dynactin’s filament are all different lengths. Here, our structure shows that the four N termini all exit the shoulder at residue 100. To accommodate the different binding site lengths, the two N termini bound to the top protofilament are pulled taut. In contrast, the two that bind the bottom protofilament have longer sections of disorder.

We previously observed the helices of p150 entering the shoulder between the arms and splitting off to enter the hook domains. We can now trace the rest of the p150 in the shoulder. After contributing a helical hairpin to the hook domain, the C-terminal 33 residues come together with p50 subunits to form the dimerization domain. Because p150 makes an intricate network of interactions with other subunits, it is difficult to imagine it existing in isolation, at least with its current conformation. Previous studies reported the isolated C terminus of p150 (residues 1050-1286) can interact with potential adaptors including RILP (28), SNX6 (29) and HPS6 (30). It is unclear, however, whether these interactions are possible when the C terminus is embedded in the shoulder.

### Dynactin’s pointed end as an interaction hub

At the pointed end, the structure of p62 was previously elusive. Studies had identified some secondary structural elements (6) and proposed that a series of cysteines formed zinc-binding motifs (31, 32). Our structure now resolves p62’s β-sandwich, central saddle domain and long helix regions. Twelve cysteines within the p62 sequence are positioned in three zinc-binding motifs. A previous study predicted that eight of these formed a cross-brace RING domain (32). We find instead that these residues form two separate folds, one a Gag-like knuckle and the second a zinc ribbon (as defined by (33)). The final zinc binding motif uses residues from the N- and C-terminal halves of p62 (Figure 3A, arrow). This motif helps stabilize the correct fold of site 2 for cargo adaptor interactions.

Currently all the identified cargo adaptors that activate dynein and dynactin for processive movement contain long coiled coils. Previous work reported the structure of three adaptors, BICD2, BICDR1 and Hook3, bound to dynein and dynactin. These structures showed that the three adaptors bind similarly to a few small sites on the filament. However they lacked the resolution to identify the contact sites on the pointed end, to which the cargo adaptors appeared to bind differently (9). Using our structure, we can now identify the residues on the pointed end used to bind the cargo adaptors. These residues are largely ionic in nature. They cluster into four interaction sites, with each site consisting of a small number of residues. Of the four sites only one, site 3, was predicted from previous modelling data (21). Site 1 is a flexible, positively-charged loop that contacts all adaptors. All three adaptors make some contacts to site 2, although the residues involved differ. Site 3 and 4 only contact a subset of adaptors. Some of the residues in these sites can undergo post-translational modification. S31 in p25, a site 4 residue that binds to BICDR1 and BICD2, is a phosphorylation site (34). In addition K302 on p62, a residue in site 2 that binds to BICD2, can be ubiquitinated (35). These modifications could directly modulate cargo adaptor binding.

Our conservation analysis shows that the adaptor binding sites on p62 and p25 are conserved between holozoa (metazoa and closely related single-celled eukaryotes excluding fungi). In contrast p27, which does not contain any contact sites, shows much less surface conservation (Figure 4E). Most of the interaction interfaces are on the front face of the pointed end, with the exception of site 4, which extends round to the end face of the p25 β-helical domain. Site 4 induces a pronounced kink in the cargo adaptors which contact it, BICD2 and BICDR1 (6, 9). The function for this kink is unclear, but it may help orient the cargo optimally for transport. Compared to the front and end faces of the pointed end, the back face is less well conserved, consistent with it making no interaction with cargo adaptors.

The adaptor interaction sites are overall less conserved in fungi and other simpler eukaryotes, with only a subset of species containing a complete pointed end (36). In Saccharomyces cerevisiae, where dynactin is not used for vesicular transport, there appear to be no genes for p62, p25 or p27 (36–38). In contrast, Aspergillus nidulans, which uses dynactin and the coiled-coil cargo adaptor HookA for dynein-mediated transport of early endosomes (39), has a complete pointed end. Strikingly, the portion of p62 comprising site 1 is not conserved in A. nidulans. The long helix region is predicted to contain different secondary structure elements. Sites 2, 3 and 4 are much better conserved. Given that HookA is the only identified adaptor in A. nidulans, and that the main coiled coil in mammalian Hook3 only uses site 3, it is intriguing that all three sites are conserved.

Our structure explains previous observations suggesting that p25 is a central component for binding cargo adaptors (13, 19). p25 is in the center of the pointed end complex, holding together p62 and p27. Furthermore p25 contains two conserved interaction sites for adaptors (site 3 and 4). In contrast, p27 is dispensable for HookA binding in A. nidulans, and for some dynactin functions in C. elegans. This is consistent with our observations that p27 does not contain conserved interaction interfaces for cargo adaptors. An exception to this pattern is the reported role of p27 in localizing dynactin to the kinetochore (13). The role for p27 in this process in unclear, but may be related to either recruitment of the kinetochore-localizing cargo adaptor Spindly or other kinetochore components such as polo-like kinase 1, which is reported to bind p27’s disordered C terminus (20).

As well as binding cargo adaptors, the pointed end has been shown to interact with the p150 projection in previous EM datasets (6, 24). Our crosslinking mass spectrometry data shows that this conformation can occur in solution and our structure shows it would prevent binding of the three cargo adaptors. This suggests that the p150 arm acts to autoinhibit dynactin, and that this must be overcome to allow cargo adaptor binding. Autoinhibition is a recurring theme in the molecular motors (25, 40–44). Dynactin autoinhibition could add another layer of regulation to this complex network. In this model, all three components of the dynein-dynactin-cargo adaptor complex are autoinhibited, with inhibition overcome by stochastic activation and binding of components to drive active complex formation for long processive transport.

## Materials and methods

### Constructs and sample preparation

The following con-structs were used: pACEBac1-HOOK31-522-SNAPf-Psc-2xStrep (9); dynein tail construct containing residues 1-1455 of the human dynein heavy chain (HC) in a pACEBac1 vector contain an N-terminal His_6_-ZZ-TEV tag and fused to pDyn2 (containing genes of human IC2C, LIC2, Tctex1, LC8 and Robl1) as described (8).

Dynactin was purified from pig brains using the large scale SP-sepharose protocol (6). C-terminal Psc-Strep tagged constructs and the dynein tail construct were expressed and purified using baculovirus as previous (9).

For dynein tail-dynactin-Hook3 (TDH) complex grids, dynein tail, dynactin and HOOK3_1522_-SNAPf were mixed in a 2:1:20 molar ratio in GF150 buffer (25 mM HEPES pH 7.2; 150 mM KCl; 1 mM MgCl_2_; 5 mM DTT; 0.1 mM ATP) and incubated on ice for 15 min. The sample was crosslinked to increase the amount of complex formed by addition of 0.0125% (v/v) glutaraldehyde (Sigma-Aldrich), at room temperature for 15 min before quenching with 200 mM Tris pH 7.4 (final concentration). The sample was gel filtered using a TSKgel G4000SWXL (TOSOH Bioscience) equilibrated in 25 mM Hepes-KOH pH 7.2, 150 mM KCl, 1 mM MgCl_2_, 0.1 mM Mg.ATP, 5 mM DTT. The TDH complex was concentrated in a 100 kDa cut-off Amicon centrifugal concentrator (Merck) at 1,500 rcf to 0.1-0.2 mg/ml, and TWEEN-20 was then added to a concentration of 0.005% (w/v). 3 μl of the TDH sample were applied to freshly glow-discharged Quantifoil R2/2 300-mesh copper grids covered with a thin carbon support. Samples were incubated on grids on a Vitrobot IV (Thermo Fisher Scientific) for 45 s and blotted for 3-4.5 s at 100% humidity and 4°C, then plunged into liquid ethane.

For dynactin grids, dynactin was crosslinked at 350 nM in GF150 buffer using 0.05% glutaraldehyde for 45 minutes at 4°C. Reactions were quenched using 100 mM Tris pH 7.4. Quantifoil R2/2 300-square-mesh copper grids were covered with a thin carbon support, and glow-discharged for 70 s at 15 mA using a Pelco easiGlow system. 3 μl of sample was then applied to the grids on a ThermoFisher Vitrobot IV at 100% humidity and 4°C, incubated for 30-40 s, blotted for 3-3.5 s, then plunged into liquid ethane.

### Cryo-EM data collection and initial data processing for TDH

Electron micrograph movies were recorded using a Titan Krios (Thermo Fisher Scientific) equipped with an energy-filtered K2 detector (Gatan) at 105,000x magnification in EFTEM mode (300 kV, 40 frames, 10 s exposure, 40 e^-^/Å^2^). For TDH, movies were acquired in super-resolution mode at the MRC Laboratory of Molecular Biology (0.58 Å/pix), or in counting mode (1.07 Å/pix) at the University of Leeds. Data were collected between 1.5-3 μm underfocus using Serial EM or EPU. 4 movies per hole were collected. Correction of inter-frame movement of each pixel and dose-weighting were performed using MotionCor2 (binning the superresolution data by 2, 5 x 5 patches, excluding first 3 frames) (21). CTF parameters were estimated using GCTF (45). Micrographs with limited CTF information or with ice contamination were removed at this stage.

For each TDH dataset, Gautomatch (http://www.mrc-lmb.cam.ac.uk/kzhang/) was used to pick particles from all micrographs (4x binned) using 2D classes from EMD-4177 as a reference. 2D classification in RELION 3.0 (spherical mask size 750 Å), combined with manual inspection of particles was used to remove ice, protein aggregates and other junk particles. Initial 3D refinement was performed using EMD-4177 as an input model, low-passed to 60 Å. Each dataset was then cleaned once using 3D classification, using the output from 3D refinement as a reference. At this point datasets collected at the LMB were merged. These data were then further refined and classified.

After this step, this dataset was merged with data from Leeds and previous data (9). Pixel sizes to match the LMB K2 datasets using Chimera (46). A b-factor of +150 Å^2^ was applied to the K2 dataset, to allow for combination of particles from different detectors (44). After this combination, data were further refined, giving a final reconstruction at 5.6 Å.

### Combination of data

Three datasets were then combined to focus on dynactin: the combined TDH dataset (above); the final particle stack from the tail-dynactin-BICDR1 structure (9); the final particle stack from the dynactin structure (6). Pixel sizes were first rescaled to match the TDR dataset (1.34 Å/pix). For TDR and TDH datasets, density for dynein and adaptors were then subtracted in two steps, each using a 4 pixel soft-edge mask. Dynein heavy chains (from residue 467 to its C terminus) and dynein light intermediate chains were first subtracted in RELION 3.0. The output particles were refined to more accurately align the remaining density. We then used a second round of signal subtraction to remove the remainder of the dynein signal and the cargo adaptor. After these steps, data could be combined. The combined dataset was then subjected to a round of global refinement, initially using a 600 Å spherical mask, then a 6 pixel soft-edge mask. This resulted in an overall dynactin reconstruction (EMD-11313) at a resolution of 3.8 Å.

### Data processing for the shoulder and pointed end

Pro-cessing for the dynactin shoulder was performed in RELION 3.0. Different masks were tried with signal subtraction to improve the density of the shoulder. All of the masks tried were created using a low-pass filter of 15 Å and 6 pixel soft edge. Masks of the shoulder components alone or including a small portion of the underlying filament did not have enough signal to align. In contrast a mask including shoulder components and five underlying filament subunits improved the density of the shoulder. This mask was optimized using focused refinement without signal subtraction. The map from this refinement was closely examined to see ordered density outside the mask, and blurred density inside the mask. Using this optimization technique we found that the Arp1 subunit A nearest the barbed end could be removed from the mask, whilst half of Arp1 subunits F and G should be included. Signal subtraction was then performed, subtracting the signal outside of this mask. This was followed by local refinement using a 6 pixel soft-edge mask. The CTF parameters were then refined. This was succeeded by a round of 3D classification without alignments. Due to the size of the particle in comparison to the box size a T value of 50 was used, with 25 classes. The best class was then locally refined using a 6 pixel soft-edge mask. This map, EMD-11314, was at an overall resolution of 3.8 Å, and was used to build the majority of the shoulder. After CTF refinement, signal subtraction also was used to focus on a map consisting of the upper paddle, upper hook and distal region of the lower arm, using a 4 pixel soft-edge mask. After signal subtraction, 3D classification was performed without alignments using a T value of 180, 16 classes, and limiting resolution to 6 Å in the expectation step. We chose the class from this containing the most ordered density, and reverted to non-subtracted shoulder particles, subjecting these particles to a local masked refinement. This resulted in a map with an overall resolution of 4.6 Å. This map, EMD-11316, was low-passed to 6 Å for modelling the upper paddle, upper hook and distal region of the lower arm.

For the pointed end, RELION 3.1 was used (Scheres 2019). This version of RELION allowed us to re-center particles on masks center-of-mass during subtraction and also to reduce the box size. Subtraction was accomplished in two stages to ensure accurate subtraction. First, half of the dynactin, including the barbed end and shoulder subunits was subtracted using a 6 pixel soft-edge mask. This was followed by local refinement using a 6 pixel soft-edge mask, to enable further subtraction using more accurate angles for the remaining half of dynactin. We then optimized the mask, to exclude more of the filament in a subsequent second round of subtraction. After this subtraction it became clear that some signal from the β-sandwich domain of p62 had been removed when subtracting the density for the adaptor in the TDR and TDH complexes. We hence went back to these complexes and repeated the signal subtraction of the adaptors using a tighter mask around the adaptors to prevent subtraction of p62 signal. After processing this new subtraction as above, the β-sandwich domain of p62 was better resolved. We subjected these particles to 3D classification with no alignments, performed with a T-value of 50, 25 classes. The best class was then locally refined using a 6 pixel soft-edge mask to an overall resolution of 4.1 Å(EMD-11315).

To examine the pointed end for TDR and TDH, the complex datasets were kept separate after the first step of dynein subtraction, rather than combining these data as above. For each complex a mask around the pointed end was created including the adaptor. Signal outside of this mask was subtracted using RELION 3.1 to leave the pointed end with adaptor attached. These maps were then subjected to a round of local refinement.

All maps were post-processed using RELION 3.1, with b-factors initially estimated automatically (47). For building, each map was processed using multiple b-factors between the estimated value and zero and low passed to different resolutions, to account for the heterogeneity in resolution in the maps. For examining the interface with adaptors, lower b-factors were used, as at high b-factors signal for the adaptors was lost. Local resolution was calculated in RELION 3.1 (48). EMDA (49) (available at https://www2.mrc-lmb.cam.ac.uk/groups/murshudov/content/emda/emda.html) was used to align shoulder and pointed end maps to the map of the whole dynactin. This avoids the information loss inherent in the resampling procedure in chimera.

### Cryo-EM data collection and initial data processing to study dynactin p150 docked conformation

Electron micrograph movies were recorded using a Titan Krios (Thermo Fisher Scientific) equipped with an energy-filtered K2 detector (Gatan) in at 105,000x magnification in EFTEM mode (300 kV, 40 frames, 10 s exposure, 40 e^-^/Å^2^. Movies were acquired in counting mode for dynactin (1.16 Å/pix). Data were collected between 1.5-3 μm underfo-cus using Serial EM or EPU. 4 images per hole were collected. Correction of inter-frame movement of each pixel and dose-weighting were performed using MotionCor2 (5×5 patches) (50). CTF parameters were estimated using GCTF. Micrographs with limited CTF information or with ice contamination were removed at this stage.

For dynactin datasets, Gautomatch (http://www.mrc-lmb.cam.ac.uk/kzhang/) was used to pick particles from all micrographs (4x binned) using 2D projections of EMD-2856 as a reference. 2D classification in RELION, combined with manual inspection of particles was used to remove ice, protein aggregates and other junk particles. Initial 3D refinement was performed using EMD-2856 as an input model, low-passed to 60 Å. Each dataset was then cleaned once using 3D classification, using the output from 3D refinement as a reference. At this point datasets were unbinned and merged. This combined dataset was aligned using 2D classification, then subjected to 3D refinement, classification and Bayesian polishing.

To focus on the p150 docked conformation, these data were then merged with the dynactin particles from our combined dataset. Signal subtraction was used to focus on the pointed end. Two rounds of 3D classification were used to separate dynactin particles containing a docked p150 arm. The first round used a spherical mask to coarsely classify particles with density for p150. The second round of classification used a mask around the p150 density. The best class from this was then locally refined.

### Model building and refinement

Building was performed in COOT (51, 52). For the shoulder, we first used real-space refine in COOT to refine the dynactin model from the previous TDR structure, PDB 6F1T (9), into our new density. Secondary structure elements were first rebuilt if necessary, and fit into density. Following this sidechains were built, starting with bulky sidechains, allowing for assignment of the subunits in the shoulder. In the lower copy we could build sidechains in the paddle and hook bundles, and the arm, excluding the middle region and the distal region of the arm. In the upper copy we could build sidechains in half of the arm bundle, including the distal region. We could also assign sidechains for the dimerization domain. In other regions, secondary structure elements were built, with loops modelled into low threshold density where possible.

For the pointed end, we used real-space refine in COOT to refine the pointed end subunits from the previous TDR model (PDB 6F1T) into our density. We could then assign and build side chains for p25 and p27. For p62, side chains for its long helix were first modelled. We then built the rest of the structure, assigning side chains in the saddle region, and part of the *β*-sandwich. Two loops were not modelled due to lack of density, indicating extreme flexibility. The first (residues 89-105) is between two adjacent β-strands. The second (residues 183-218) was the disordered loop following the long helix in p62.

Model refinements were performed in REFMAC5 (53, 54). The entire dynactin model was first refined into the overall dynactin map to best fit the model into the density. For further refinement, the model was then split into three sub-models corresponding to each of the two rigid bodies used for masked refinement of the shoulder and pointed end, and the remainder of the filament. These filament subunits were refined into the overall map. Refinement proceeded on these three sub-models separately, iterating between manual optimization of model geometry using coot real-space refine and automated real-space refinement in REFMAC5. These refinements consisted of 20 iterations using a refinement weighting of 0.0001, with hydrogen atoms included. The early refinements on the filament imposed non-crystallographic symmetry onto the 7 barbed-end proximal Arp1 subunits (Chain ID A-G). In the first refinements of both the pointed end and the shoulder, the maps were initially low-pass filtered to 6 Å in order to fit the model into the areas with lower resolution. The high-resolution maps were then used for subsequent refinements, in which the side chain conformations were optimized. For each sub-model, refinements continued until the model validation scores stopped improving, as calculated using PHENIX validation tools (55, 56). Boundaries between sub-models were refined in PHENIX, using restraints on other parts of the model.

To assess pointed end interactions with cargo adaptors/p150 arm, we first used rigid-body fitting to place our new pointed end model into the appropriate map: EMD-11317 for pointed end-BICDR1 interactions; EMD-11318 for pointed end-Hook3 interactions; EMD-2860 for pointed end-BICD2 interactions (6); EMD-11319 for pointed end-p150 interactions. We used rigid-body fitting to place previous structures of TDB (PDB 6F3A), TDR (PDB 6F1T), TDH (PDB 6F38) or dynactin with the p150 (PDB 5ADX) into the appropriate map, then removed components other than the cargo adaptor/p150 arm. For the pointed end-BICDR1 and pointed-BICD2 models, cargo adaptors were refined using PHENIX (at 6 Å and 8 Å respectively), to resolve clashes with the pointed end. For the pointed end-Hook3 model, the second coiled-coil was modelled into density. For the p150 arm, the registry of the coiled coils was updated to be consistent with our mass spectrometry crosslinking data. We then examined the density connecting our new pointed end structure with the cargo adaptor/p150 arm to discern likely interacting residues. Contact sites were only considered where there is density for both dynactins pointed end and cargo adaptor, with connecting density. Solvent accessible surface areas were approximated using Pymol.

### Cross-linking mass spectrometry

200 μg dynactin at 3 μM in GF150 buffer was crosslinked with 1.5 mM BS3 for 2 hours at 4°C. The reaction was then quenched using 160 mM Tris pH 7.4. The crosslinked samples were cold-acetone precipitated and resuspended in 8 M urea and 100mM NH_4_HCO_3_. Proteins were reduced with 10mM DTT and alkylated with 50mM iodoacetamide. Following alkylation, proteins were digested with Lys-C (Pierce) at an enzyme-to-substrate ratio of 1:100 for 4 h at 22°C and, after diluting the urea to 1.5 M with 100mM NH_4_HCO_3_ solution, and further digested with trypsin (Pierce) at an enzyme-to-substrate ratio of 1:20.

Digested peptides were eluted from StageTips and split into two for parallel crosslink enrichment by strong cation-exchange chromatography (SCX) and size exclusion chromatography (SEC), and were dried in a vacuum concentrator (Eppendorf). For SCX, eluted peptides were dissolved in mobile phase A (30% acetonitrile (v/v), 10 mM KH2PO4, pH 3) before strong cation exchange chromatography (100 Œ 2.1 mm Poly Sulfoethyl A column; Poly LC). The separation of the digest used a gradient into mobile phase B (30% acetonitrile (v/v), 10 mM KH_2_PO_4_, pH 3, 1 M KCl) at a flow rate of 200 ţl min1. Ten 1-minute fractions in the high-salt range were collected and cleaned by StageTips, eluted and dried for subsequent liquid chromatography with tandem mass spectrometry (LC-MS/MS) analysis. For peptideSEC, peptides were fractionated on an ÄKTA Pure system (GE Healthcare) using a Superdex Peptide 3.2/300 (GE Healthcare) at a flow rate of 10 ţl min1 using 30% (v/v) acetonitrile and 0.1% (v/v) trifluoroacetic acid as mobile phase. Five 50-ţl fractions were collected and dried for subsequent LC-MS/MS analysis.

Samples for analysis were resuspended in 0.1% v/v formic acid, 1.6% v/v acetonitrile. LC-MS/MS analysis was conducted in duplicate for SEC fractions and triplicate for SCX fractions, performed on an Orbitrap Fusion Lumos Tribrid mass spectrometer (Thermo Fisher Scientific) coupled on-line with an Ultimate 3000 RSLCnano system (Dionex, Thermo Fisher Scientific). The sample was separated and ionized by a 50 cm EASY-Spray column (Thermo Fisher Scientific). Mobile phase A consisted of 0.1% (v/v) formic acid and mobile phase B of 80% v/v acetonitrile with 0.1% v/v formic acid. Flow-rate of 0.3 μl min1 using gradients optimized for each chromatographic fraction from offline fractionation ranging from 2% mobile phase B to 45% mobile phase B over 90 min, followed by a linear increase to 55% and 95% mobile phase B in 2.5 min, respectively. The MS data were acquired in data-dependent mode using the top-speed setting with a three second cycle time. For every cycle, the full scan mass spectrum was recorded in the Orbitrap at a resolution of 120,000 in the range of 400 to 1,600 m/z. Ions with a precursor charge state between 3+ and 7+ were isolated and fragmented. Fragmentation by higher-energy collisional dissociation (HCD) employed a decision tree logic with optimized collision energies (57). The fragmentation spectra were then recorded in the Orbitrap with a resolution of 30,000. Dynamic exclusion was enabled with single repeat count and 60-s exclusion duration.

A recalibration of the precursor m/z was conducted based on high-confidence (< 1% false discovery rate (FDR)) linear peptide identifications. The recalibrated peak lists were searched against the sequences and the reversed sequences (as decoys) of crosslinked peptides using the Xi software suite (v.1.6.745) for identification (58). The following parameters were applied for the search: MS1 accuracy = 3 ppm; MS2 accuracy = 10 ppm; enzyme = trypsin (with full tryptic specificity) allowing up to four missed cleavages; crosslinker = BS3 with an assumed reaction specificity for lysine, serine, threonine, tyrosine and protein N termini; fixed modifications = carbamidomethylation on cysteine; variable modifications = oxidation on methionine, hydrolyzed/aminolyzed BS3 from reaction with ammonia or water on a free crosslinker end. The identified candidates were filtered to 2% FDR on link level using XiFDR v.1.1.26.58 (59).

Crosslinks were then plotted on the structure in Pymol and their lengths were calculated. Disordered loops were modelled in Coot, to assess instances where at least one crosslinked residue was in a loop. The p150 projection, the flexible N and C termini of Arp11, and the disordered loop in p62 were not modelled, due to their length. Crosslinks under 30 Å (C_*α*_-C_*α*_distance) were considered valid. Crosslinked residue pairs over 30 Å apart were examined to see if small changes in the conformation of dynactin could enable valid crosslinks to form. For crosslinks between subunits where multiple copies exist in dynactin (e.g. Arp1), the shortest crosslink was assessed.

### Bioinformatics

PSI-BLAST and JACKHMMER profile-based sequence searches were used to identity eukaryotic homologs of p25, p27 and p62. Towards this, we provided the sequences of p25, p27 and p62 from Sus scrofa as initial query inputs to search against UniProt/TrEMBL databases (with an e-value cut-off = 0.001). Further, more distant homologs of p25, p27 and p62 of Sus scrofa were identified by providing respective multiple sequence alignments of the first set of above-identified homologs as queries to JACKHMMER to search against UniProt/TrEMBL (evalue = 0.001). Sequences from a diverse set of eukaryotes were aligned using MAFFT (60). Weblogo was used to visualize these alignments for the interacting residues (61). ConSurf was used to calculate per-residue conservation scores from these alignments (62). To visualize surface conservation, conservation scores were rendered on the models of the pointed end. Side chain conservation was then rendered onto the density for dynactins pointed end. The standard ConSurf color-blind-friendly color scheme was used for visualization.

### Figure rendering and formatting

Density images for figures were rendered using ChimeraX (63) or Chimera v1.13.1 (64), and model-only images were rendered using Pymol (DeLano 2002). Formatting for Biorxiv used a modified LaTeX template from the Henriques Lab (https://www.overleaf.com/latex/templates/henriqueslab-biorxiv-template/nyprsybwffws#.Wp8hF1Cnx-E)

## Data availability

Cryo-EM maps are available at the EMDB (EMD-11313, EMD-11314, EMD-11315, EMD-11316, EMD-11317, EMD-11318, EMD-11319). Model coordinates are deposited at the PDB (6ZNL, 7ZNM, 6ZNN, 6ZNO, 6ZO4). Crosslinking mass spectrometry data are deposited at PRIDE (PXD020084).

## Author contributions

C.K.L. and A.P.C. conceived the research. C.K.L performed the cryo-EM work, and expressed and purified proteins. F.J.O. conducted the cross-linking mass spectrometry experiments. B.S. collated and aligned the pointed end protein sequences. C.K.L and S.E.L built and refined the dynactin model. A.P.C. and C.L. prepared the manuscript. All authors edited the manuscript.

## ACKNOWLEDGEMENTS

We thank L. Urnavicius and K. Zhang for helpful discussion and datasets for TDR and dynactin; T. Nakane, S. Scheres, A. Murzin and S. Shakeel for helpful discussions; J. Grimmett and T. Darling for scientific computing support; and F. Abid Ali and S. Chaaban for comments on the manuscript. We acknowledge the MRC – Laboratory of Molecular Biology Electron Microscopy Facility for access and support of electron microscopy sample preparation and data collection; the University of Leeds Electron Microscopy facility for help with data collection and Diamond for access and support of the Cryo-EM facilities at the UK national electron bio-imaging centre (eBIC), proposal EM18086, funded by the Wellcome Trust, MRC and BBSRC. This work was funded by grants from the Wellcome Trust (WT210711) and the Medical Research Council, UK (MC_UP_A025_1011) to A.P.C., Wellcome Trust Senior Research Fellowship (103139) to J.R, Deutsche Forschungsgemeinschaft project no. 426290502. We acknowledge

---

**Supplementary Figure 1.**
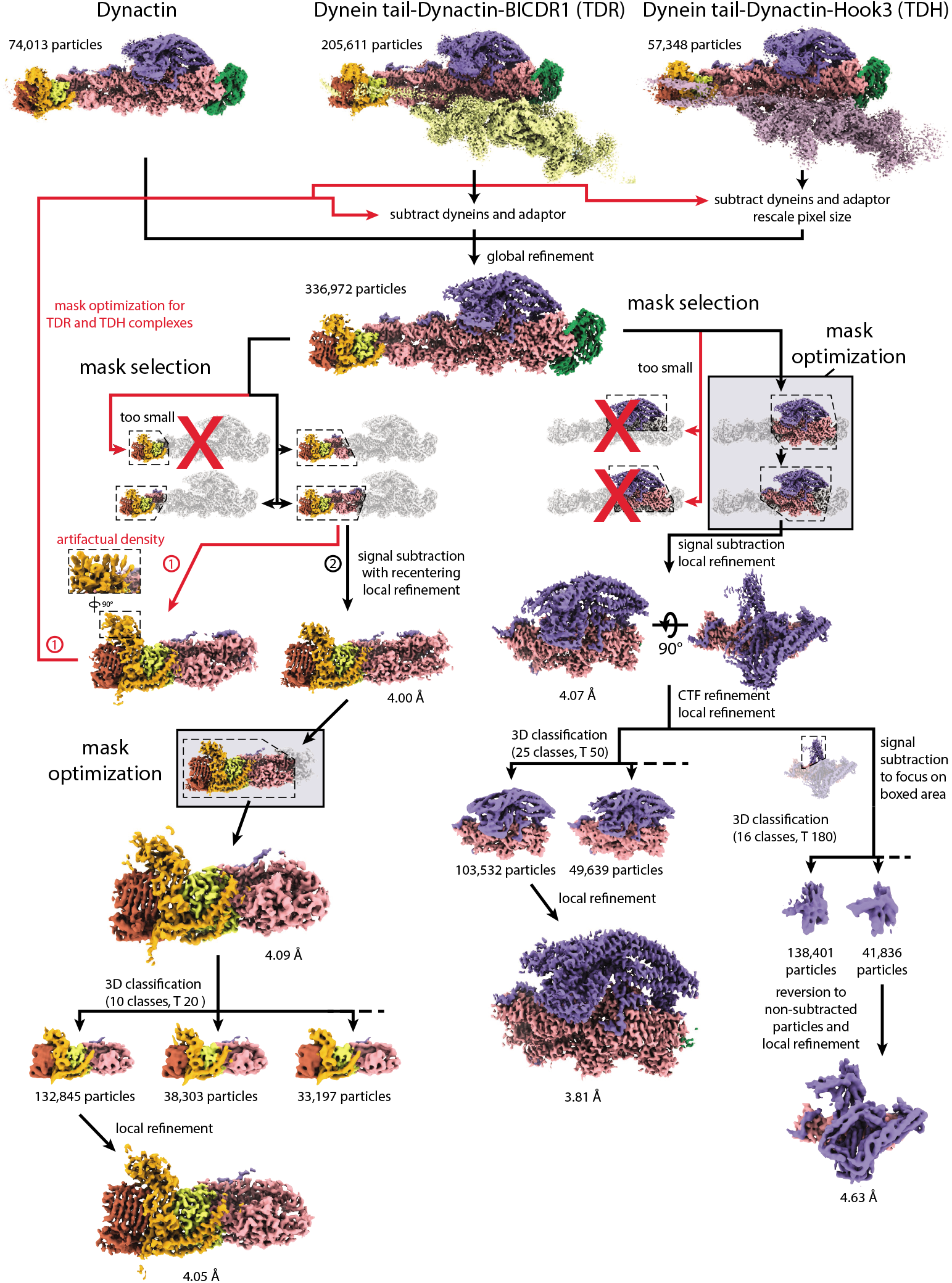
Data processing pipeline for high resolution dynactin maps. Flowchart showing processing steps for the production of high resolution dynactin maps. Mask optimization steps are highlighted in boxes. For the pointed end, the first attempt resulting in artefactual density is labelled in red (1), with the second successful attempt shown in black (2).

**Supplementary Figure 2.**
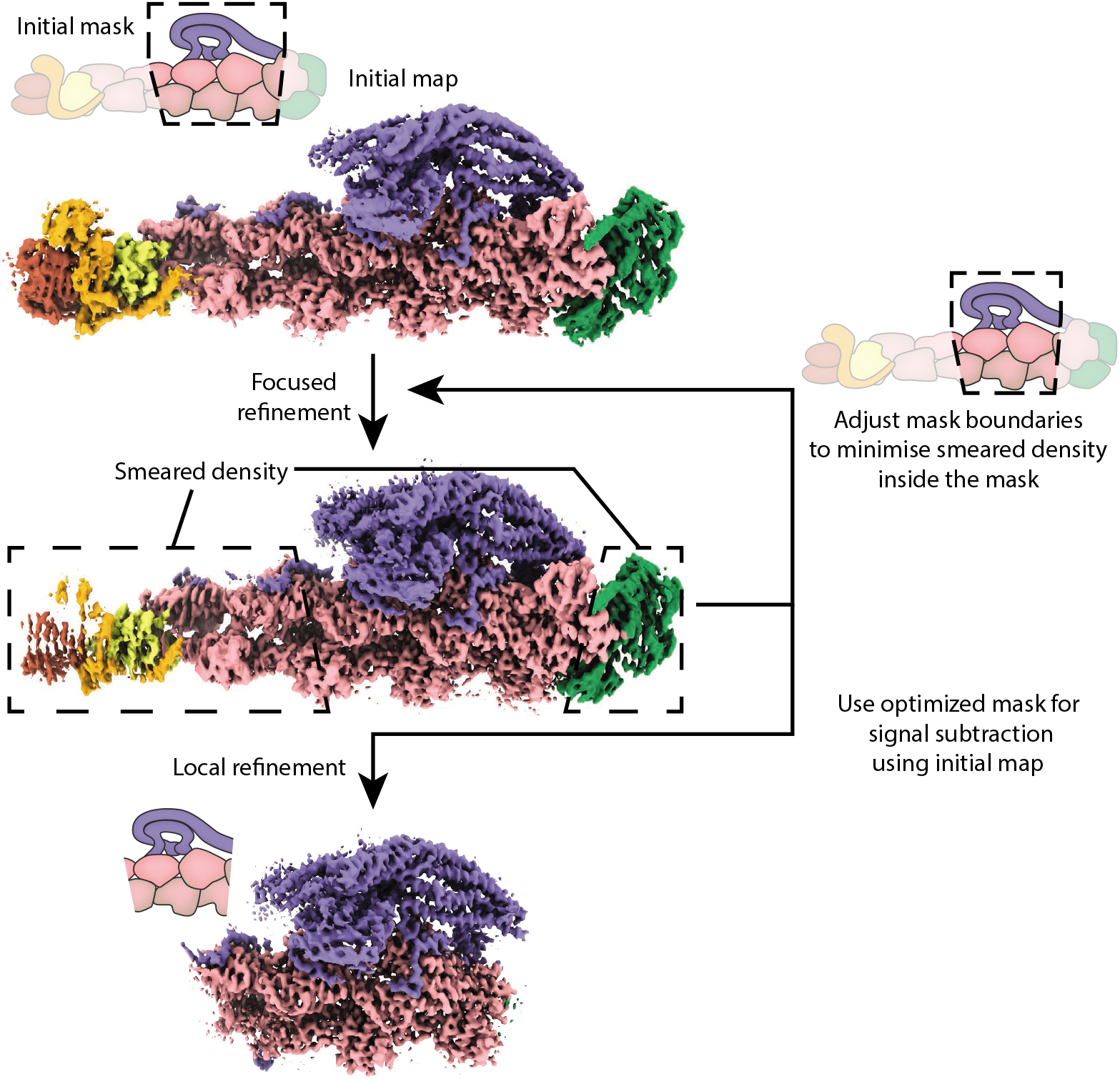
Mask optimization for signal subtraction. Flowchart detailing the steps of mask optimization for signal subtraction.

**Supplementary Figure 3.**
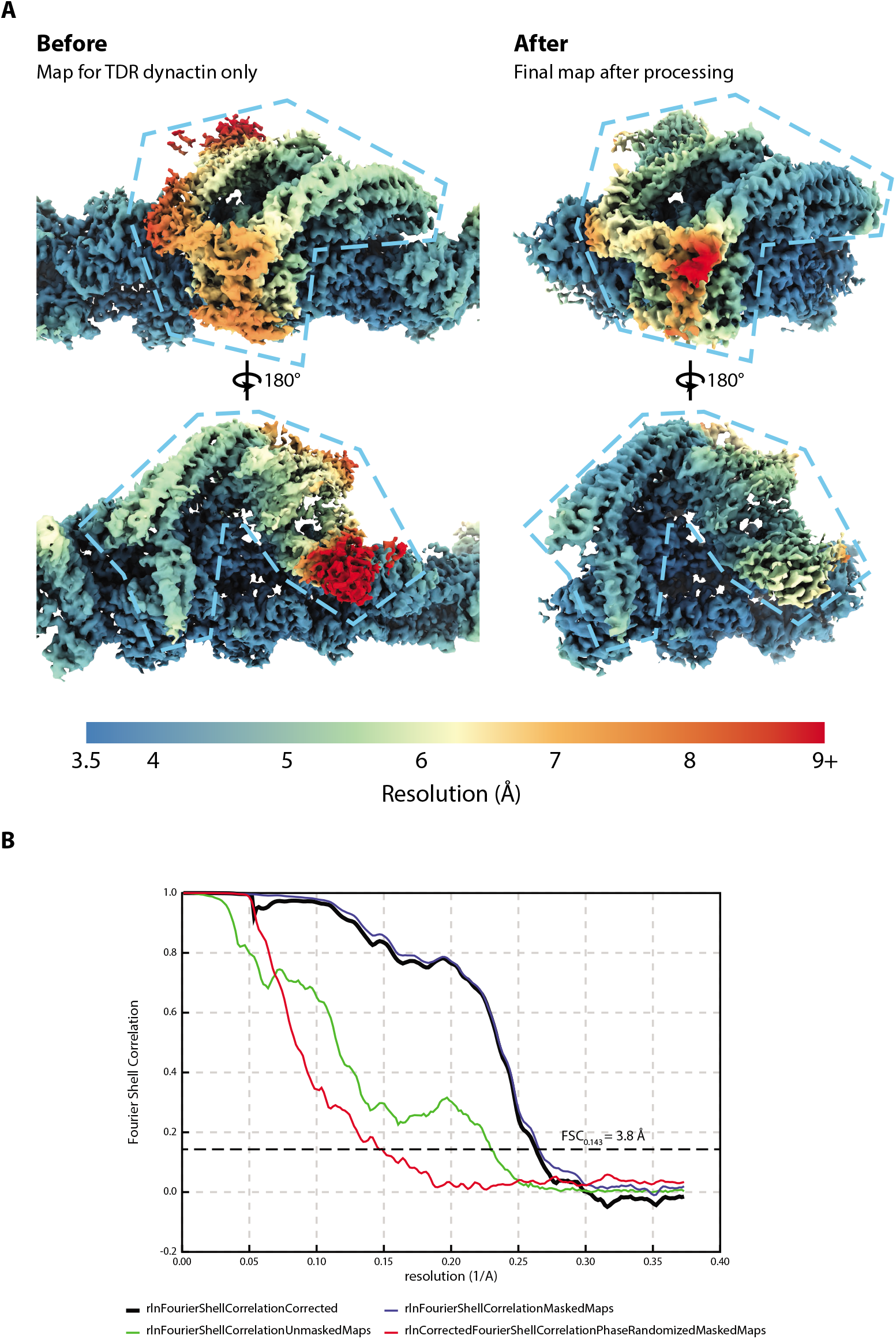
Local resolution of the shoulder map. **A.** Local resolution (as calculated by RELION) for the shoulder region of dynactin to show the improvement in density quality. The before map shows dynactin density from the dynein tail-dynactin-BICDR1 (TDR, left) and the after map shows the final shoulder map (right). Dynactins shoulder is marked by a dotted line. **B.** Gold-standard FSC curve for the new shoulder map, showing map resolution at the FSC0.143 cutoff.

**Supplementary Figure 4:**
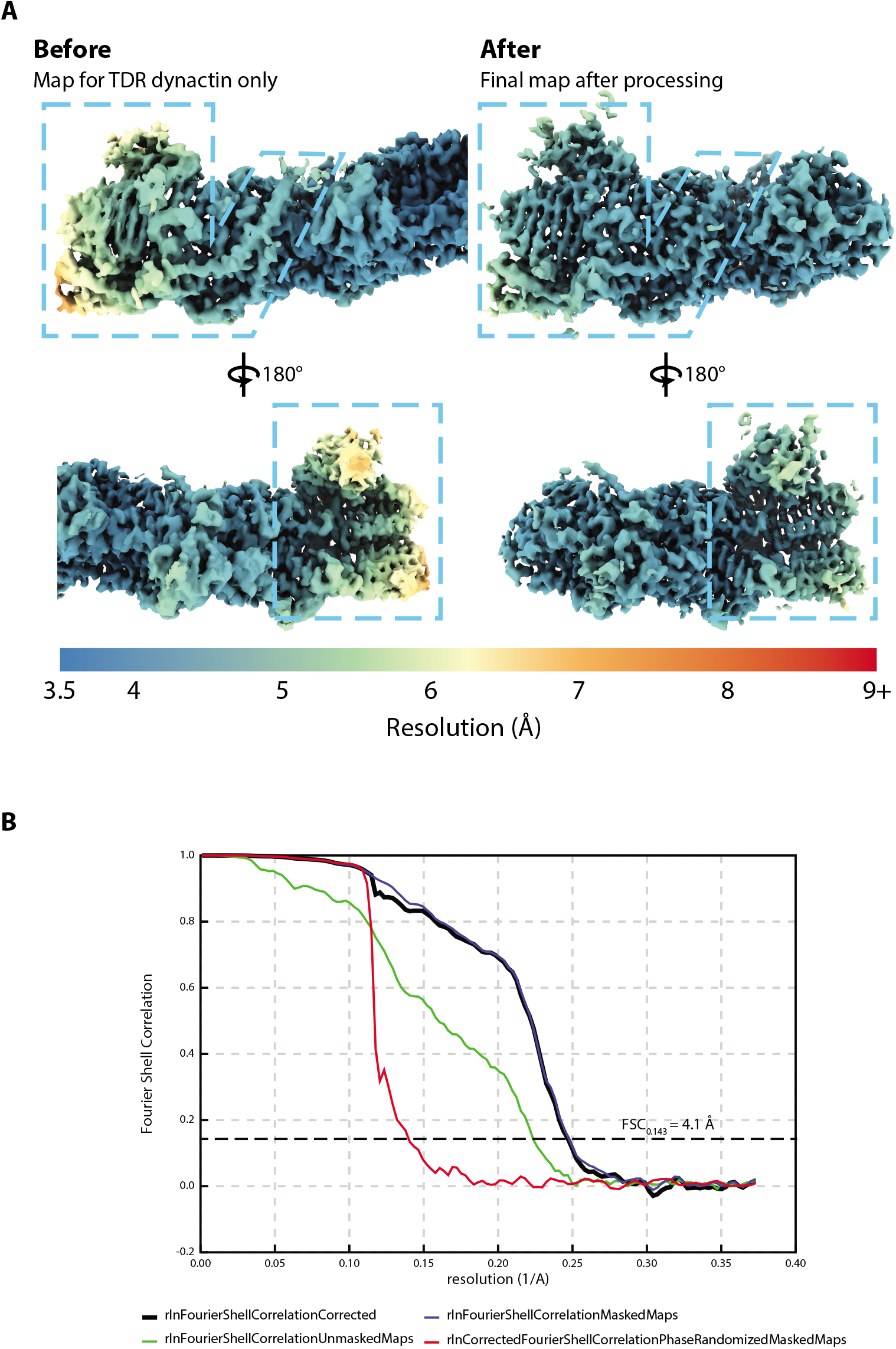
Local resolution for pointed end map. **A.** Local resolution (as calculated by RELION) for the pointed end region of dynactin. The before map shows dynactin density from the dynein tail-dynactin-BICDR1 (TDR, left) and the after map shows the final pointed end map (right). Dynactins pointed end is marked by a dotted line. **B.** Gold-standard FSC curve for the new pointed end map, showing map resolution at the FSC0.143 cutoff.

**Supplementary Figure 5.**
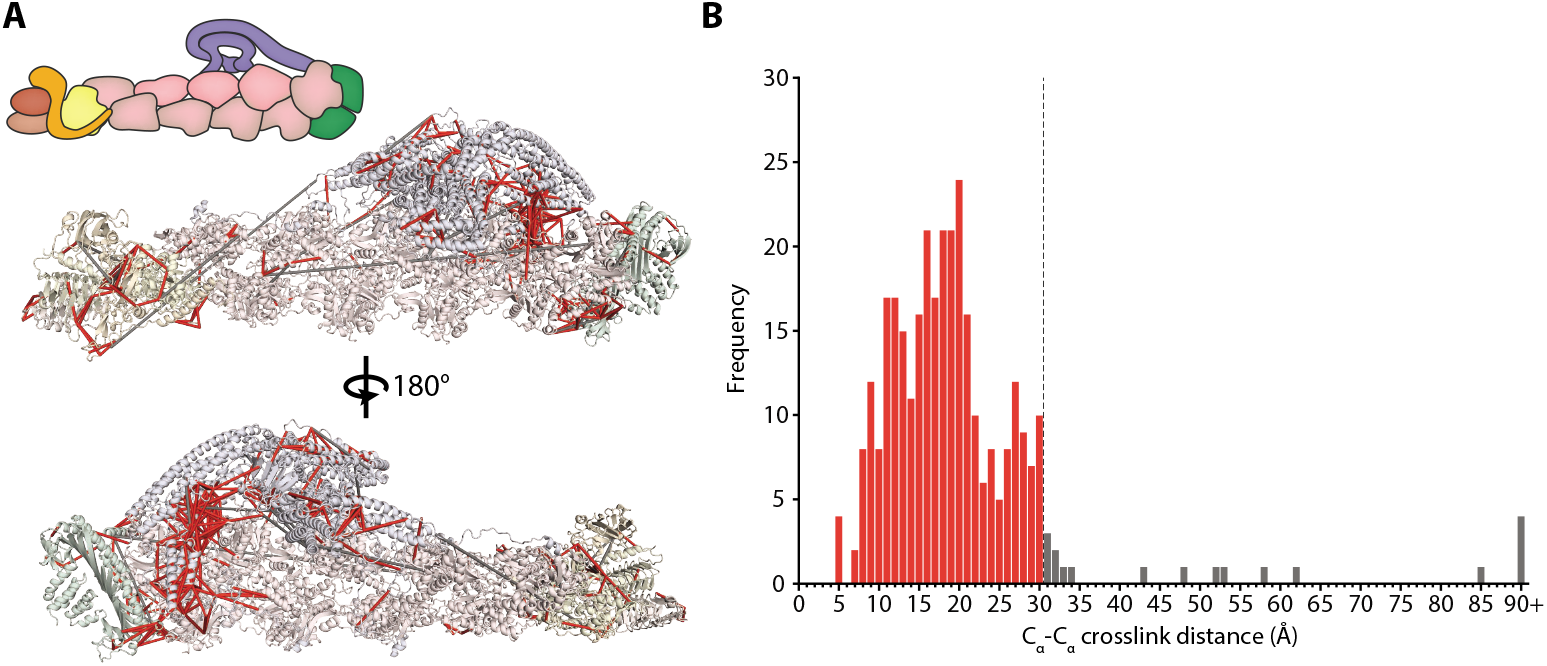
Crosslinking mass spectrometry of dynactin. **A.** Crosslinks (lines) plotted onto the dynactin structure (pastel cartoon). Valid crosslinks, under 30 Å in length (C_*α*_-C_*α*_), are shown in red, whereas overlength crosslinks are shown in gray. **B.** Histogram of C_*α*_-C_*α*_ distances between crosslinked residue pairs, with the 30 Å cutoff marked using a dotted line. Valid crosslinks are shown in red, whereas overlength crosslinks are shown in gray.

**Supplementary Figure 6.**
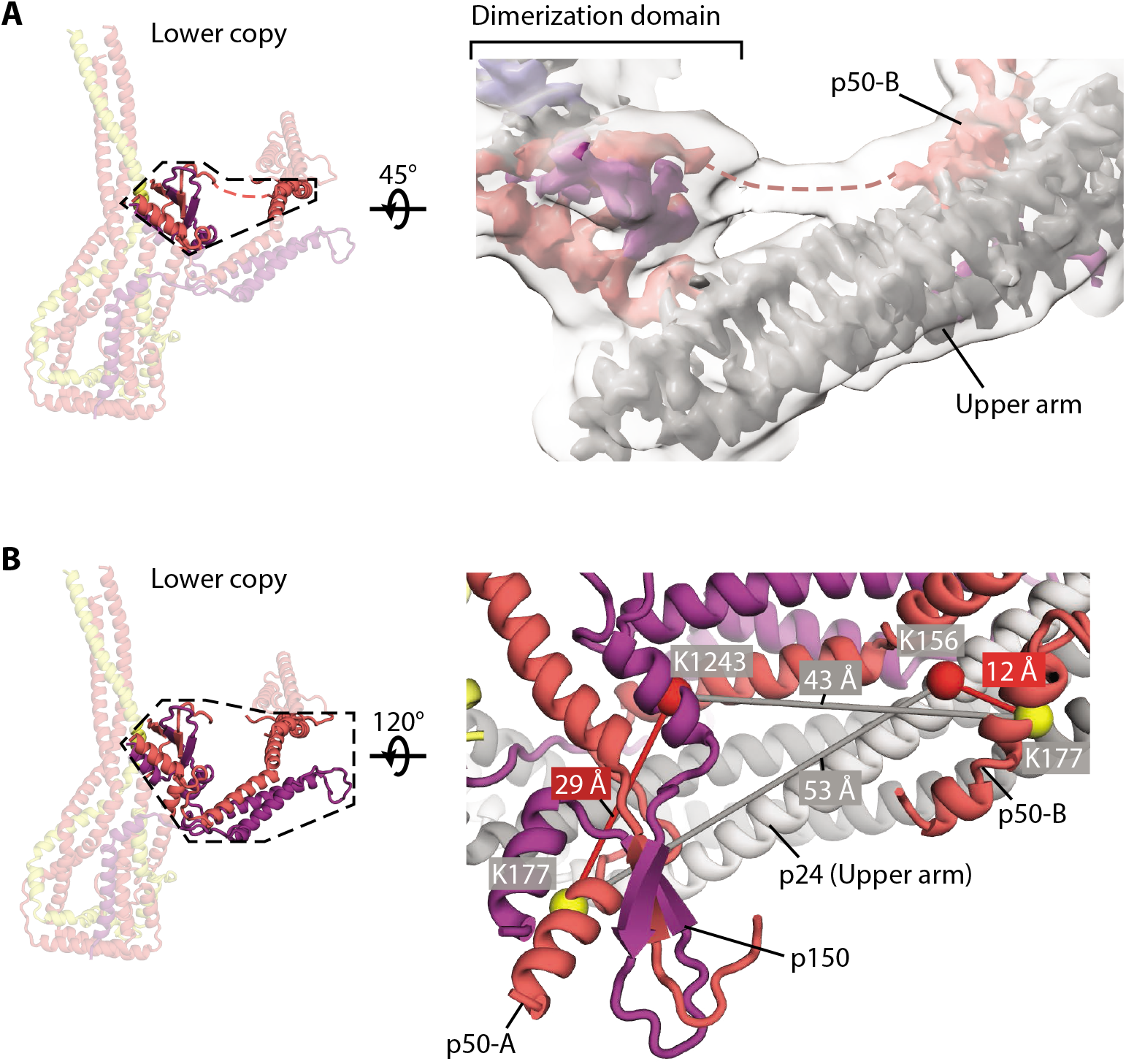
Asymmetry of the p50 subunits within the shoulder. **A.** Diagram showing the connection in p50-B between residue 186 and residue 205 (dotted line). To show the connection the high resolution map (colored density) was filtered to 6 Å and blurred using a b factor of +100 Å^2^ (transparent density). **B.** Model diagram showing two examples of key validating crosslinks (<30 Å C_*α*_-C_*α*_distance, red lines) for the asymmetric organization of p50 subunits A and B. These two crosslinks are between p50 residue K177 (red cartoon, yellow spheres) and the upper arm p150 residue K1243 (purple cartoon, red sphere), and p24 residue K156 (gray cartoon, red sphere). p50-A K177 satisfies the crosslink with p150 K1243, while not satisfying the crosslink with p24 K156 (gray). p50-B K177 on the other hand satisfies the crosslink with p24 K156, but not with p150 K1243 (gray).

**Supplementary Figure 7:**
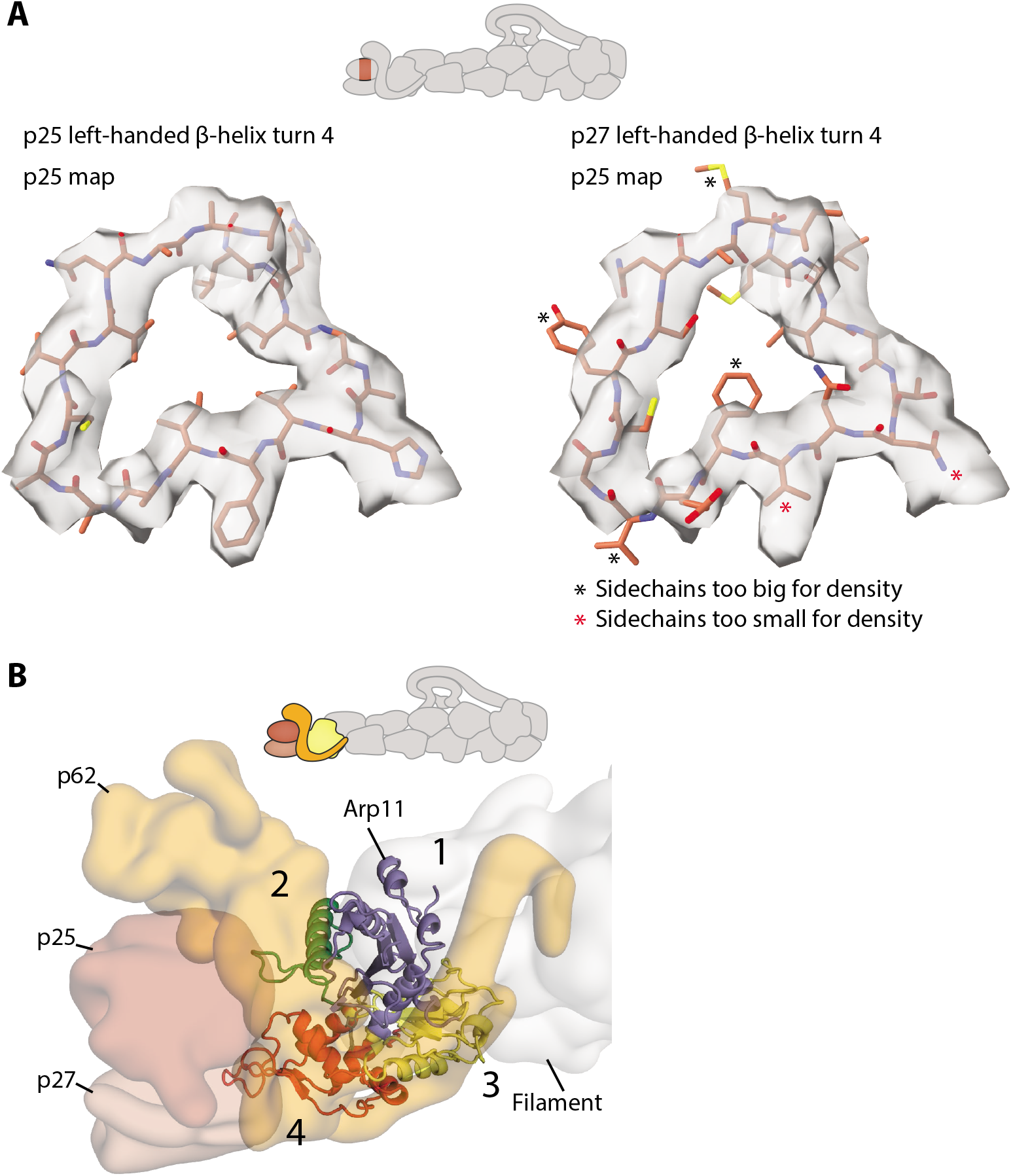
Features at the pointed end. **A.** Electron density from p25 at the pointed end showing that p25, but not p27, fit the density. Equivalent residues from turn 4 of the left-handed β-helix were taken from p25 (left, residues 84-103) and p27 (right, residues 81-100), and fit into the p25 density. This shows that sidechains from p25 fit well, whereas those in p27 do not. Sidechains in p27 that do not fit the density are marked by asterisks. **B.** Subdomain diagram of Arp11, showing contacts with different pointed end subunits. Arp domains are colored by subdomain, with p62 (orange), p25 (brown) and p27 (light brown) in transparent surface.

**Supplementary Figure 8:**
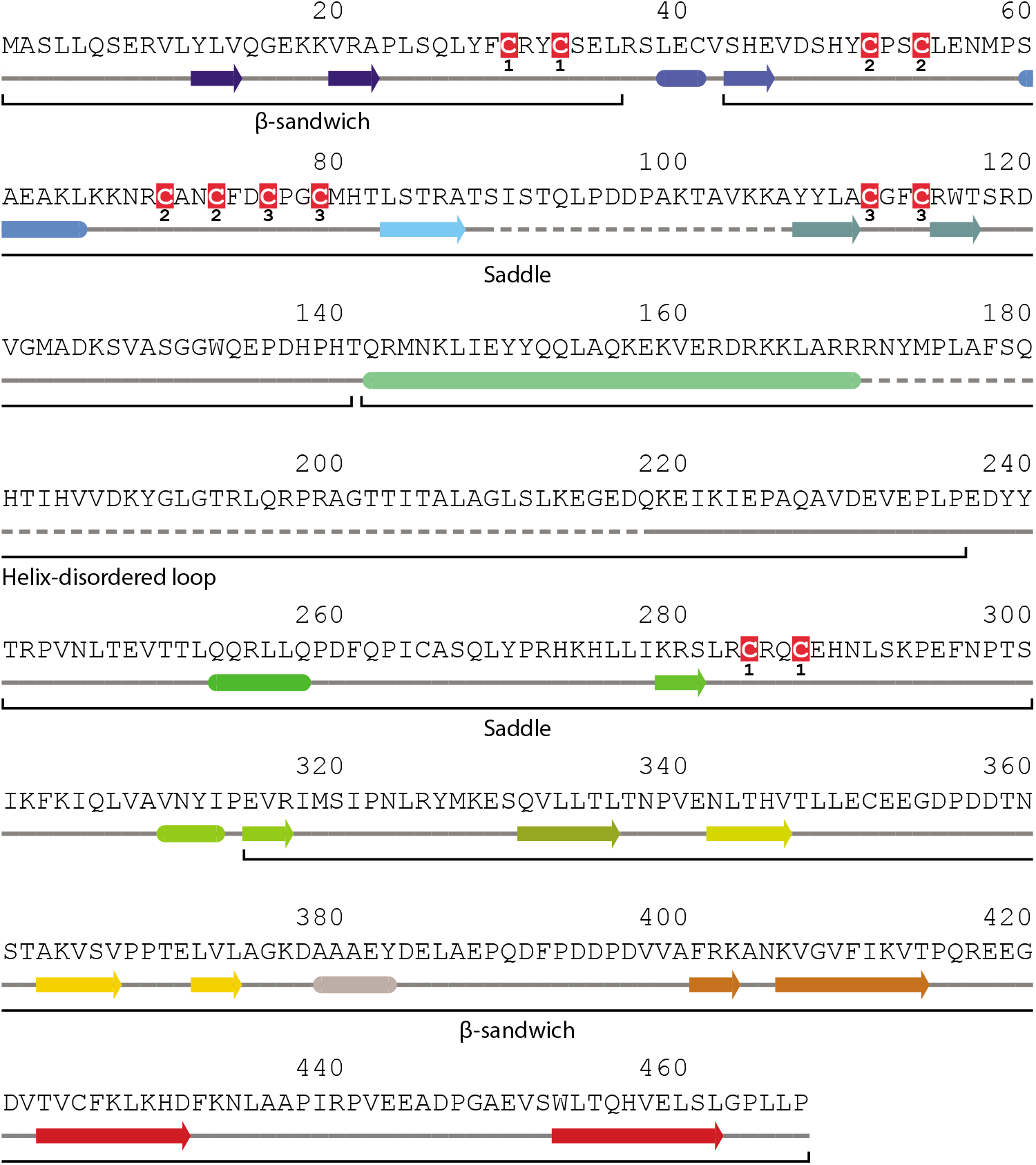
The secondary structure of p62. Secondary structure diagram of p62, with secondary structure elements colored as in Figure 3B. The three different regions of p62 (β-sandwich, saddle and helix-disordered loop) are marked, to denote the parts of sequence that contribute to each one. Cysteines are highlighted and labelled to show the residues belonging to each of the three metal-binding motifs.

**Supplementary Figure 9:**
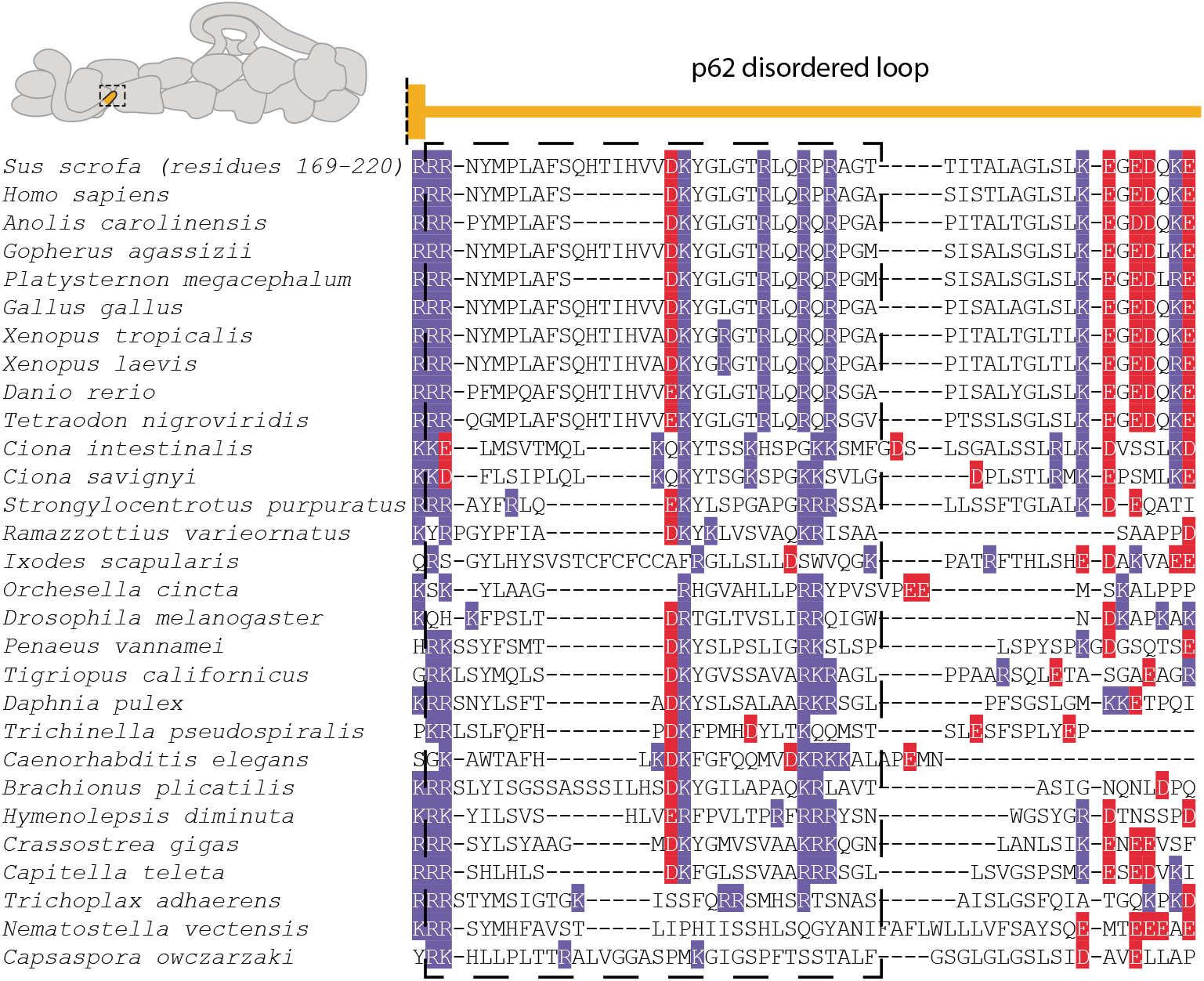
p62 interaction site 1 sequence alignment. Sequence alignments for the p62 disordered loop (site 1) from diverse eukaryotes. The end of the long helix preceding the loop is shown above. Positively- and negatively-charged residues are highlighted in blue and red respectively. This first half of the loop, which is positively-charged in all organisms aligned, is highlighted in a dotted box.

**Supplementary Figure 10.**
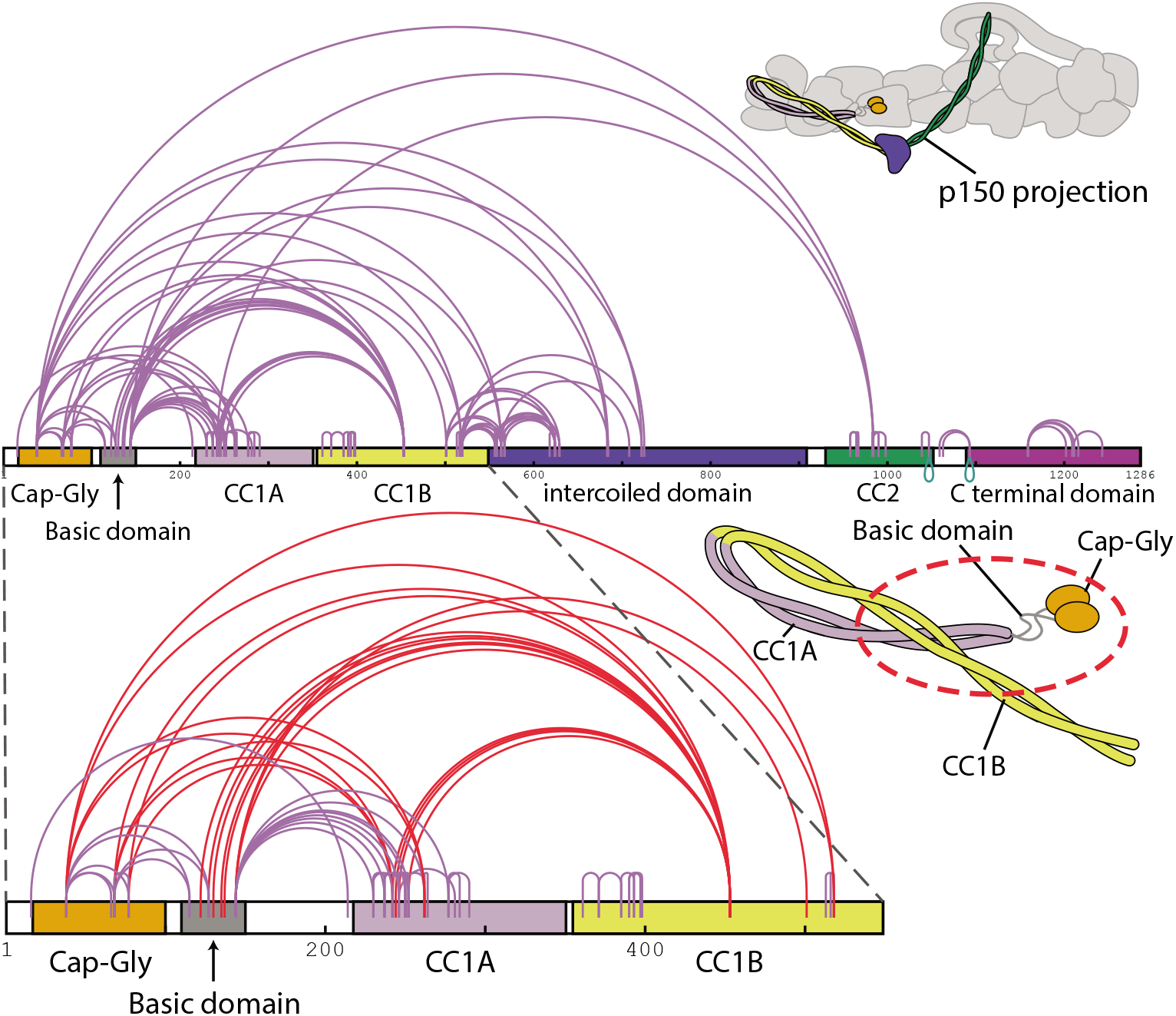
Crosslinking mass spectrometry within the p150 projection of dynactin. Diagram showing all crosslinks identified within p150 by crosslinking mass spectrometry. Subdomains of p150 are colored and labelled on the schematic and accompanying cartoon. Crosslinks are marked by purple lines. The inset highlights the crosslinks between CC1B and other regions of the N terminus of p150, supporting its antiparallel arrangement (red crosslinks).

## Bibliography

1. Samara L Reck-Peterson, William B Redwine, Ronald D Vale, and Andrew P Carter. The cytoplasmic dynein transport machinery and its many cargoes. Nature reviews. Molecular cell biology, 19(6):382–398, 2018. ISSN 1471-0080. doi: 10.1038/s41580-018-0004-3.

2. D A Schafer, S R Gill, J A Cooper, J E Heuser, and T A Schroer. Ultrastructural analysis of the dynactin complex: an actin-related protein is a component of a filament that resembles F-actin. The Journal of Cell Biology, 126(2):403–412, jul 1994. ISSN 0021-9525. doi: 10.1083/jcb.126.2.403.

3. D M Eckley, S R Gill, K A Melkonian, J B Bingham, H V Goodson, J E Heuser, and T A Schroer. Analysis of dynactin subcomplexes reveals a novel actin-related protein associated with the arp1 minifilament pointed end. The Journal of cell biology, 147(2):307–20, oct 1999. ISSN 0021-9525. doi: 10.1083/jcb.147.2.307.

4. Hiroshi Imai, Akihiro Narita, Yuichiro Maéda, and Trina A Schroer. Dynactin 3D structure: implications for assembly and dynein binding. Journal of molecular biology, 426(19): 3262–3271, sep 2014. ISSN 1089-8638. doi: 10.1016/j.jmb.2014.07.010.

5. Saikat Chowdhury, Stephanie A Ketcham, Trina A Schroer, and Gabriel C Lander. Structural organization of the dynein-dynactin complex bound to microtubules. Nature Structural & Molecular Biology, 22(4):345–347, mar 2015. ISSN 1545-9993. doi: 10.1038/nsmb.2996.

6. L. Urnavicius, K. Zhang, A. G. Diamant, C. Motz, M. A. Schlager, M. Yu, N. A. Patel, C. V. Robinson, and A. P. Carter. The structure of the dynactin complex and its interaction with dynein. Science, 347(6229):1441–1446, 2015. ISSN 0036-8075. doi: 10.1126/science.aaa4080.

7. R J McKenney, W Huynh, M E Tanenbaum, G Bhabha, and R D Vale. Activation of cytoplasmic dynein motility by dynactin-cargo adapter complexes. Science, 345(6194):337–341, 2014. doi: 10.1126/science.1254198.

8. Max A Schlager, Ha Thi Hoang, Linas Urnavicius, Simon L Bullock, and Andrew P Carter. In vitro reconstitution of a highly processive recombinant human dynein complex. EMBO Journal, 33(17):1855–1868, 2014. doi: 10.15252/embj.201488792.

9. Linas Urnavicius, Clinton K. Lau, Mohamed M. Elshenawy, Edgar Morales-Rios, Carina Motz, Ahmet Yildiz, and Andrew P. Carter. Cryo-EM shows how dynactin recruits two dyneins for faster movement. Nature, 554(7691):202–206, feb 2018. ISSN 0028-0836. doi: 10.1038/nature25462.

10. S Karki and E L Holzbaur. Affinity chromatography demonstrates a direct binding between cytoplasmic dynein and the dynactin complex. The Journal of biological chemistry, 270(48): 28806–11, dec 1995. ISSN 0021-9258. doi: 10.1074/jbc.270.48.28806.

11. KT Vaughan and R B Vallee. Cytoplasmic dynein binds dynactin through a direct interaction between the intermediate chains and p150Glued. The Journal of cell biology, 131(6 Pt 1): 1507–16, dec 1995. ISSN 0021-9525. doi: 10.1083/jcb.131.6.1507.

12. Courtney M Schroeder, Jonathan M L Ostrem, Nicholas T Hertz, and Ronald D Vale. A Ras-like domain in the light intermediate chain bridges the dynein motor to a cargo-binding region. eLife, 3, oct 2014. ISSN 2050-084X. doi: 10.7554/eLife.03351.

13. José B. Gama, Cláudia Pereira, Patrícia A. Simões, Ricardo Celestino, Rita M. Reis, Daniel J. Barbosa, Helena R. Pires, Cátia Carvalho, João Amorim, Ana X. Carvalho, Dhanya K. Cheerambathur, and Reto Gassmann. Molecular mechanism of dynein recruitment to kinetochores by the RodZw10Zwilch complex and Spindly. The Journal of Cell Biology, 216(4):943–960, apr 2017. ISSN 0021-9525. doi: 10.1083/jcb.201610108.

14. In-Gyun Lee, Mara A. Olenick, Malgorzata Boczkowska, Clara Franzini-Armstrong, Erika L. F. Holzbaur, and Roberto Dominguez. A conserved interaction of the dynein light intermediate chain with dynein-dynactin effectors necessary for processivity. Nature Communications, 9(1):986, dec 2018. ISSN 2041-1723. doi: 10.1038/s41467-018-03412-8.

15. Karin A Melkonian, Kerstin C Maier, Jamie E Godfrey, Michael Rodgers, and Trina A Schroer. Mechanism of dynamitin-mediated disruption of dynactin. The Journal of biological chemistry, 282(27):19355–64, jul 2007. ISSN 0021-9258. doi: 10.1074/jbc.M700003200.

16. Frances Ka Yan Cheong, Lijuan Feng, Ali Sarkeshik, John R. Yates, and Trina A. Schroer. Dynactin integrity depends upon direct binding of dynamitin to Arp1. Molecular Biology of the Cell, 25(14):2171–2180, jul 2014. ISSN 1059-1524. doi: 10.1091/mbc.e14-03-0842.

17. Jun Zhang, Xuanli Yao, Lauren Fischer, Juan F. Abenza, Miguel A. Peñalva, and Xin Xiang. The p25 subunit of the dynactin complex is required for dyneinearly endosome interaction. The Journal of Cell Biology, 193(7):1245–1255, jun 2011. ISSN 1540-8140. doi: 10.1083/jcb.201011022.

18. Ting-Yu Yeh, Nicholas J Quintyne, Brett R Scipioni, D Mark Eckley, and Trina A Schroer. Dynactin’s pointed-end complex is a cargo-targeting module. Molecular biology of the cell, 23(19):3827–37, oct 2012. ISSN 1939-4586. doi: 10.1091/mbc.E12-07-0496.

19. Rongde Qiu, Jun Zhang, and Xin Xiang. No Title. 293(40), oct 2018. ISSN 1083-351X.

20. Ting-Yu Yeh, Anna K Kowalska, Brett R Scipioni, Frances Ka Yan Cheong, Meiying Zheng, Urszula Derewenda, Zygmunt S Derewenda, and Trina A Schroer. Dynactin helps target Polo-like kinase 1 to kinetochores via its left-handed beta-helical p27 subunit. The EMBO Journal, 32(7):1023–1035, mar 2013. ISSN 0261-4189. doi: 10.1038/emboj.2013.30.

21. Wenjun Zheng. Probing the Energetics of Dynactin Filament Assembly and the Binding of Cargo Adaptor Proteins Using Molecular Dynamics Simulation and Electrostatics-Based Structural Modeling. Biochemistry, 56(1):313–323, jan 2017. ISSN 0006-2960. doi: 10.1021/acs.biochem.6b01002.

22. Xiao-chen C Bai, Eeson Rajendra, Guanghui Yang, Yigong Shi, and Sjors HW Scheres. Sampling the conformational space of the catalytic subunit of human gamma-secretase. eLife, 4, dec 2015. ISSN 2050-084X. doi: 10.7554/eLife.11182.

23. Sjors Scheres. Single-particle processing in RELION-3.1, 2019.

24. Kei Saito, Takashi Murayama, Tomone Hata, Takuya Kobayashi, Keitaro Shibata, Saiko Kazuno, Tsutomu Fujimura, Takashi Sakurai, and Yoko Y Toyoshima. Conformational diversity of dynactin sidearm and domain organization of its subunit p150. Molecular biology of the cell, page mbcE20010031, 2020. ISSN 1939-4586. doi: 10.1091/mbc.E20-01-0031.

25. Suvranta K. Tripathy, Sarah J. Weil, Chen Chen, Preetha Anand, Richard B. Vallee, and Steven P. Gross. Autoregulatory mechanism for dynactin control of processive and diffusive dynein transport. Nature Cell Biology, 16(12):1192–1201, dec 2014. ISSN 1465-7392. doi: 10.1038/ncb3063.

26. C M Schroeder and R D Vale. Assembly and activation of dynein-dynactin by the cargo adaptor protein Hook3. Journal of Cell Biology, 214(3):309–318, 2016. doi: 10.1083/jcb.201604002.

27. Kei Saito, Takashi Murayama, Tomone Hata, Takuya Kobayashi, Keitaro Shibata, Saiko Kazuno, Tsutomu Fujimura, Takashi Sakurai, and Yoko Y. Toyoshima. Domain organization and conformational change of dynactin p150. bioRxiv, page 459040, dec 2018. doi: 10.1101/459040.

28. Marie Johansson, Nuno Rocha, Wilbert Zwart, Ingrid Jordens, Lennert Janssen, Coenraad Kuijl, Vesa M Olkkonen, and Jacques Neefjes. Activation of endosomal dynein motors by stepwise assembly of Rab7-RILP-p150Glued, ORP1L, and the receptor betalll spectrin. The Journal of cell biology, 176(4):459–471, 2007. doi: papers3://publication/doi/10.1083/jcb.200606077.

29. Zhi Hong, Yanrui Yang, Cheng Zhang, Yang Niu, Ke Li, Xi Zhao, and Jia-Jia Liu. The retromer component SNX6 interacts with dynactin p150Glued and mediates endosome-to-TGN transport. Cell Research, 19(12):1334–1349, dec 2009. ISSN 1001-0602. doi: 10.1038/cr.2009.130.

30. Ke Li, Lin Yang, Cheng Zhang, Yang Niu, Wei Li, and Jia-Jia Liu. HPS6 interacts with dynactin p150Glued to mediate retrograde trafficking and maturation of lysosomes. Journal of cell science, 127(Pt 21):4574–88, nov 2014. ISSN 1477-9137. doi: 10.1242/jcs.141978.

31. Jorge A. Garces, Imran B. Clark, David I. Meyer, and Richard B. Vallee. Interaction of the p62 subunit of dynactin with Arp1 and the cortical actin cytoskeleton. Current Biology, 9 (24):1497–1502, dec 1999. ISSN 0960-9822. doi: 10.1016/S0960-9822(00)80122-0.

32. S Karki, M K Tokito, and E L Holzbaur. A dynactin subunit with a highly conserved cysteine-rich motif interacts directly with Arp1. The Journal of biological chemistry, 275(7):4834–9, feb 2000. ISSN 0021-9258. doi: 10.1074/jbc.275.7.4834.

33. S. S. Krishna, Indraneel Majumdar, and Nick V. Grishin. Structural classification of zinc fingers: SURVEY AND SUMMARY. Nucleic Acids Research, 31(2):532–550, jan 2003. ISSN 13624962. doi: 10.1093/nar/gkg161.

34. Philipp Mertins, Feng Yang, Tao Liu, D R Mani, Vladislav A Petyuk, Michael A Gillette, Karl R Clauser, Jana W Qiao, Marina A Gritsenko, Ronald J Moore, Douglas A Levine, Reid Townsend, Petra Erdmann-Gilmore, Jacqueline E Snider, Sherri R Davies, Kelly V Ruggles, David Fenyo, R Thomas Kitchens, Shunqiang Li, Narciso Olvera, Fanny Dao, Henry Rodriguez, Daniel W Chan, Daniel Liebler, Forest White, Karin D Rodland, Gordon B Mills, Richard D Smith, Amanda G Paulovich, Matthew Ellis, and Steven A Carr. Ischemia in tumors induces early and sustained phosphorylation changes in stress kinase pathways but does not affect global protein levels. Molecular & cellular proteomics: MCP, 13(7): 1690–704, jul 2014. ISSN 1535-9484. doi: 10.1074/mcp.M113.036392.

35. Vyacheslav Akimov, Inigo Barrio-Hernandez, Sten V F Hansen, Philip Hallenborg, Anna-Kathrine Pedersen, Dorte B Bekker-Jensen, Michele Puglia, Stine D K Christensen, Jens T Vanselow, Mogens M Nielsen, Irina Kratchmarova, Christian D Kelstrup, Jesper V Olsen, and Blagoy Blagoev. UbiSite approach for comprehensive mapping of lysine and N-terminal ubiquitination sites. Nature structural & molecular biology, 25(7):631–640, 2018. ISSN 1545-9985. doi: 10.1038/s41594-018-0084-y.

36. Björn Hammesfahr and Martin Kollmar. Evolution of the eukaryotic dynactin complex, the activator of cytoplasmic dynein. BMC Evolutionary Biology, 12(1):95, jun 2012. ISSN 1471-2148. doi: 10.1186/1471-2148-12-95.

37. Elaine Yeh, Robert V. Skibbens, Judy W. Cheng, E. D. Salmon, and Kerry Bloom. Spindle dynamics and cell cycle regulation of dynein in the budding yeast, Saccharomyces cerevisiae. Journal of Cell Biology, 130(3):687–700, 1995. ISSN 00219525. doi: 10.1083/jcb.130.3.687.

38. J K Moore, M D Stuchell-Brereton, and J A Cooper. Function of dynein in budding yeast: mitotic spindle positioning in a polarized cell. Cell Motility and the Cytoskeleton, 66(8): 546–555, 2009. doi: 10.1002/cm.20364.

39. Jun Zhang, Rongde Qiu, Herbert N. Arst, Miguel A. Peñalva, and Xin Xiang. HookA is a novel dyneinearly endosome linker critical for cargo movement in vivo. The Journal of Cell Biology, 204(6), 2014.

40. L A Amos. Brain dynein crossbridges microtubules into bundles. Journal of Cell Science, 93 (Pt 1):19–28, 1989.

41. Kristen J Verhey and Jennetta W Hammond. Traffic control: regulation of kinesin motors. Nature Reviews: Molecular Cell Biology, 10(11):765–777, nov 2009. ISSN 1471-0080. doi: 10.1038/nrm2782.

42. Takayuki Torisawa, Muneyoshi Ichikawa, Akane Furuta, Kei Saito, Kazuhiro Oiwa, Hiroaki Kojima, Yoko Y. Toyoshima, and Ken’ya Furuta. Autoinhibition and cooperative activation mechanisms of cytoplasmic dynein. Nature Cell Biology, 16(11):1118–1124, sep 2014. ISSN 1465-7392. doi: 10.1038/ncb3048.

43. Shin-ichi Terawaki, Asuka Yoshikane, Yoshiki Higuchi, and Kaori Wakamatsu. Structural basis for cargo binding and autoinhibition of Bicaudal-D1 by a parallel coiled-coil with homotypic registry. Biochemical and biophysical research communications, 460(2):451–6, may 2015. ISSN 1090-2104. doi: 10.1016/j.bbrc.2015.03.054.

44. Kai Zhang, Helen E. Foster, Arnaud Rondelet, Samuel E. Lacey, Nadia Bahi-Buisson, Alexander W. Bird, and Andrew P. Carter. Cryo-EM Reveals How Human Cytoplasmic Dynein Is Auto-inhibited and Activated. Cell, 169(7):1303–1314, 2017. ISSN 10974172. doi: 10.1016/j.cell.2017.05.025.

45. Kai Zhang, X-C Xiao-chen Bai, A Brown, IS Fernandez, E Hanssen, M Condron, YH Tan, J Baum, SHW Scheres, and X-C Xiao-chen Bai. Gctf: Real-time CTF determination and correction. J Struct Biol, 193(1):1–12, jan 2016. ISSN 1095-8657. doi: 10.1016/j.jsb.2015.11.003.

46. Max E. Wilkinson, Ananthanarayanan Kumar, and Ana Casañal. Methods for merging data sets in electron cryo-microscopy. Acta Crystallographica Section D Structural Biology, 75 (9):782–791, sep 2019. ISSN 2059-7983. doi: 10.1107/S2059798319010519.

47. Peter B. Rosenthal and Richard Henderson. Optimal Determination of Particle Orientation, Absolute Hand, and Contrast Loss in Single-particle Electron Cryomicroscopy. Journal of MolecularBiology, 333(4):721–745, oct 2003. ISSN 0022-2836. doi: 10.1016/J.JMB.2003.07.013.

48. Alp Kucukelbir, Fred J Sigworth, and Hemant D Tagare. Quantifying the local resolution of cryo-EM density maps. Nature Methods, 11(1):63–65, nov 2014. ISSN 1548-7091. doi: 10.1038/nmeth.2727.

49. Rangana Warshamanage and Garib N Murshudov. EMDA: Electron Microscopy Difference and Average map tools (Version 1.1.2)., 2020.

50. Jasenko Zivanov, Takanori Nakane, Björn O Forsberg, Dari Kimanius, Wim JH Hagen, Erik Lindahl, and Sjors HW Scheres. New tools for automated high-resolution cryo-EM structure determination in RELION-3. eLife, 7, nov 2018. ISSN 2050-084X. doi: 10.7554/eLife.42166.

51. P Emsley and K Cowtan. Coot: model-building tools for molecular graphics. Acta Crystallogr D Biol Crystallogr, 60(Pt 12 Pt 1):2126–2132, 2004. doi: 10.1107/S0907444904019158.

52. P Emsley, B Lohkamp, W G Scott, and K Cowtan. Features and development of Coot. Acta Crystallogr D Biol Crystallogr, 66(Pt 4):486–501, apr 2010. ISSN 1399-0047. doi: 10.1107/S0907444910007493.

53. G N Murshudov, A A Vagin, and E J Dodson. Refinement of macromolecular structures by the maximum-likelihood method. Acta crystallographica. Section D, Biological crystallography, 53(Pt 3):240–55, may 1997. ISSN 0907-4449. doi: 10.1107/S0907444996012255.

54. Robert A Nicholls, Michal Tykac, Oleg Kovalevskiy, and Garib N Murshudov. Current approaches for the fitting and refinement of atomic models into cryo-EM maps using CCP-EM. Acta crystallographica. Section D, Structural biology, 74(Pt 6):492–505, jun 2018. ISSN 2059-7983. doi: 10.1107/S2059798318007313.

55. Pavel V. Afonine, Bruno P. Klaholz, Nigel W. Moriarty, Billy K. Poon, Oleg V. Sobolev, Thomas C. Terwilliger, Paul D. Adams, Alexandre Urzhumtsev, and IUCr. New tools for the analysis and validation of cryo-EM maps and atomic models. Acta Crystallographica Section D Structural Biology, 74(9):814–840, sep 2018. ISSN 2059-7983. doi: 10.1107/S2059798318009324.

56. Christopher J. Williams, Jeffrey J. Headd, Nigel W. Moriarty, Michael G. Prisant, Lizbeth L. Videau, Lindsay N. Deis, Vishal Verma, Daniel A. Keedy, Bradley J. Hintze, Vincent B. Chen, Swati Jain, Steven M. Lewis, W. Bryan Arendall, Jack Snoeyink, Paul D. Adams, Simon C. Lovell, Jane S. Richardson, and David C. Richardson. MolProbity: More and better reference data for improved all-atom structure validation. Protein Science, 27(1): 293–315, jan 2018. ISSN 09618368. doi: 10.1002/pro.3330.

57. Lars Kolbowski, Marta L. Mendes, and Juri Rappsilber. Optimizing the Parameters Governing the Fragmentation of Cross-Linked Peptides in a Tribrid Mass Spectrometer. Analytical Chemistry, 89(10):5311–5318, may 2017. ISSN 0003-2700. doi: 10.1021/acs.analchem.6b04935.

58. Marta L Mendes, Lutz Fischer, Zhuo A Chen, Marta Barbon, Francis J O’Reilly, Sven H Giese, Michael BohlkeSchneider, Adam Belsom, Therese Dau, Colin W Combe, Martin Graham, Markus R Eisele, Wolfgang Baumeister, Christian Speck, and Juri Rappsilber. An integrated workflow for crosslinking mass spectrometry. Molecular Systems Biology, 15(9), sep 2019. ISSN 1744-4292. doi: 10.15252/msb.20198994.

59. Lutz Fischer and Juri Rappsilber. Quirks of Error Estimation in Cross-Linking/Mass Spectrometry. Analytical Chemistry, 89(7):3829–3833, apr 2017. ISSN 0003-2700. doi: 10.1021/acs.analchem.6b03745.

60. Fábio Madeira, Young Mi Park, Joon Lee, Nicola Buso, Tamer Gur, Nandana Madhusoodanan, Prasad Basutkar, Adrian R.N. Tivey, Simon C. Potter, Robert D. Finn, and Rodrigo Lopez. The EMBL-EBI search and sequence analysis tools APIs in 2019. Nucleic Acids Research, 47(W1):W636–W641, jul 2019. ISSN 13624962. doi: 10.1093/nar/gkz268.

61. Gavin E Crooks, Gary Hon, John-Marc Chandonia, and Steven E Brenner. WebLogo: a sequence logo generator. Genome research, 14(6):1188–90, jun 2004. ISSN 1088-9051. doi: 10.1101/gr.849004.

62. Haim Ashkenazy, Shiran Abadi, Eric Martz, Ofer Chay, Itay Mayrose, Tal Pupko, and Nir Ben-Tal. ConSurf 2016: an improved methodology to estimate and visualize evolutionary conservation in macromolecules. Nucleic Acids Research, 44(Web Server issue):W344, 2016. ISSN 1362-4962. doi: 10.1093/NAR/GKW408.

63. Thomas D. Goddard, Conrad C. Huang, Elaine C. Meng, Eric F. Pettersen, Gregory S. Couch, John H. Morris, and Thomas E. Ferrin. UCSF ChimeraX: Meeting modern challenges in visualization and analysis. Protein Science, 27(1):14–25, jan 2018. ISSN 09618368. doi: 10.1002/pro.3235.

64. E F Pettersen, T D Goddard, C C Huang, G S Couch, D M Greenblatt, E C Meng, and T E Ferrin. UCSF Chimera-a visualization system for exploratory research and analysis. J Comput Chem, 25(13):1605–1612, 2004. doi: 10.1002/jcc.20084.

